# The junctional mechanosensor AmotL2 regulates YAP promotor accessibility

**DOI:** 10.1101/2023.01.13.523596

**Authors:** Aarren J. Mannion, Honglei Zhao, Yuanyuan Zhang, Ylva von Wright, Otto Bergman, Hanna M. Björck, Pipsa Saharinen, Lars Holmgren

## Abstract

Endothelial cells (ECs) are constantly exposed to mechanical forces in the form of fluid shear stress, extracellular stiffness, and cyclic strain. How these forces are sensed by ECs remains an understudied aspect in the homeostatic regulation of the circulatory system. Angiomotin-like 2 (AmotL2) is localised to EC junctions and is required for alignment and actin reorganisation under conditions of high shear stress. Here we show that AmotL2 crucially regulates transcription and promotor activity of the YAP gene. Functionally, density-dependent proliferation of ECs *in vitro* and proliferation of a subpopulation of ECs within the inner aortic arch, were both reliant on AmotL2 and Yap/Taz endothelial expression *in vivo*. Mechanistically, depletion of AmotL2 led to altered nuclear morphology, chromatin accessibility and suppression of YAP-promotor activity through increased H3K27me3 mediated by the polycromb repressive complex component EZH2. Our data describe a previously unknown role for junctional mechanotransduction in shaping the epigenetic landscape and transcriptional regulation of YAP in vascular homeostasis.

## Introduction

The endothelium, the inner most layer of blood vessels, is constantly exposed to the mechanical force exerted by blood flow. How endothelial cells (EC) sense this mechanical force and respond is context dependent and plays an important role in both vascular homeostasis and the development of disease. The laminar flow of larger blood vessels such as the descending aorta maintains a homeostatic anti-inflammatory and anti-proliferative state, whereas regions of the vasculature that are exposed to atherogenic or disturbed flow such as bifurcations, branch points, and the inner arch of the aorta are more prone to inflammation, EC proliferation and the development of disease^1,2^.

At the cell-cell contacts of ECs, junctional molecules sense mechanical force and transduce these signals intracellularly, allowing adaptation of the endothelium to changes in mechanical stimuli. Key examples of vascular junctional mechanosensors include the VEGFR2, VE-cadherin, and PECAM1^3^ junctional complex and more recently, PLEXIND1 which sense fluid shear stress in a conformation dependent manner in endothelial cells and shape their response to such forces^4^. Forces acting on junctional complexes are transduced by activation of intracellular secondary signalling molecules and mobilisation of the cytoskeletal network. Recently, the junctional molecule AmotL2 was shown to regulate endothelial and nuclear morphology in endothelial cells of the aorta, through regulation of the actin cytoskeleton and formation of the junctional complex between VE-cadherin and p120^5^. The importance of this junctional complex in correctly sensing and transducing fluid shear stress is exemplified by the development of vascular diseases such as the formation of abdominal aortic aneurysm in mice deleted for endothelial AmotL2^5^.

Yes-associated protein (YAP) and its paralogue TAZ, are transcriptional coactivators and key transducers of mechanical force^6^. Much focus has been given to the localization and activity of YAP, and its role as a regulator of proliferation and apoptosis and more recently in shaping chromatin accessibility^7^. However, little is known about the transcriptional or epigenetic control of the *YAP* gene.

Here we uncover a feedback loop of regulation between AmotL2 and YAP, where YAP activity is critical in driving AmotL2 expression within the aortic endothelium and that in turn, AmotL2 is indispensable for endothelial *YAP* transcription and promotor accessibility through modulation of histone activity. Our results reveal that mechanical forces sensed at cell-cell junctions by AmotL2 directly influence global chromatin accessibility and impact on EZH2 activity leading to modulation of promotor activity of the mechanosensor *YAP*. To our knowledge, we believe this is the first description of junctional dependent epigenetic regulation of *YAP* which controls vascular homeostasis.

## Results

### Endothelial AmotL2 is regulated by YAP *in vitro*

Previous work has shown that AmotL2 expression is driven by YAP in multiple cellular contexts^8,9^, and so we set out to confirm this in an endothelial setting. We performed RNA interference (RNAi) to knockdown *YAP* by lentiviral shRNA in Human arterial endothelial cells (HUAEC) which reduced the expression of *YAP* mRNA along with a concurrent reduction in *AmotL2* as well as *CTGF* and *ANKRD1*, two well defined YAP target genes (Fig1a). Individual and co-depletion of *YAP* and *TAZ* significantly reduced *AmotL2* expression (SuppFig1a), however, co-depletion of both *YAP* and *TAZ* did not have any additive effect in reducing *AmotL2* levels (SuppFig1a). Individual and co-depletion of *YAP* and *TAZ* by siRNA did not reduce other YAP target genes in HUAEC cells, however, a significant reduction in YAP/TAZ target genes were observed when siRNA depletion of *YAP* and/or *TAZ* was employed in Human umbilical vein endothelial cells (HUVEC), along with a reduction in *AmotL2* mRNA (SuppFig1b), which was also apparent when using shRNA (SuppFig1c). Analysis of protein expression showed that YAP shRNA led to a concurrent reduction of AmotL2 in both HUAEC (Fig1b) and HUVEC (SuppFig1d). Conversely, the overexpression of a constitutively active YAP mutant (5SA) led to increased AmotL2 expression in HUVEC (SuppFig1e-f). To analyse whether stiffness of the microenvironment was able to increase *AmotL2* expression through increased YAP activity, HUVEC were plated to soft and stiff hydrogels and RNA isolated. Results indicated that stiffer matrices upregulated *AmotL2* transcripts, compared to softer hydrogels (SuppFig1g). To verify that YAP binds to the *AmotL2* promotor to initiate transcription, we analysed existing ChIP-seq data using the ChIP-Atlas database (https://chip-atlas.org/). Screening of *AmotL2* transcriptional start site (TSS) for both YAP and TEAD1 binding indicated enrichment across 3 different cell lines (SuppFig2a), with overlapping enrichment of YAP, TEAD1, and TEAD4 (SuppFig2b). Importantly, YAP and TEAD1 and 4 were also enriched at the promotors of *CTGF* and *CYR61* (SuppFig2c). Predicted target genes of YAP indicated *AmotL2* as the top hit in the ChIP-Atlas database at 1, 5 and 10 Kb from the TSS (SuppFig2d). Taken together, these results confirm that YAP transcriptionally drives the expression of AmotL2.

**Fig 1.**
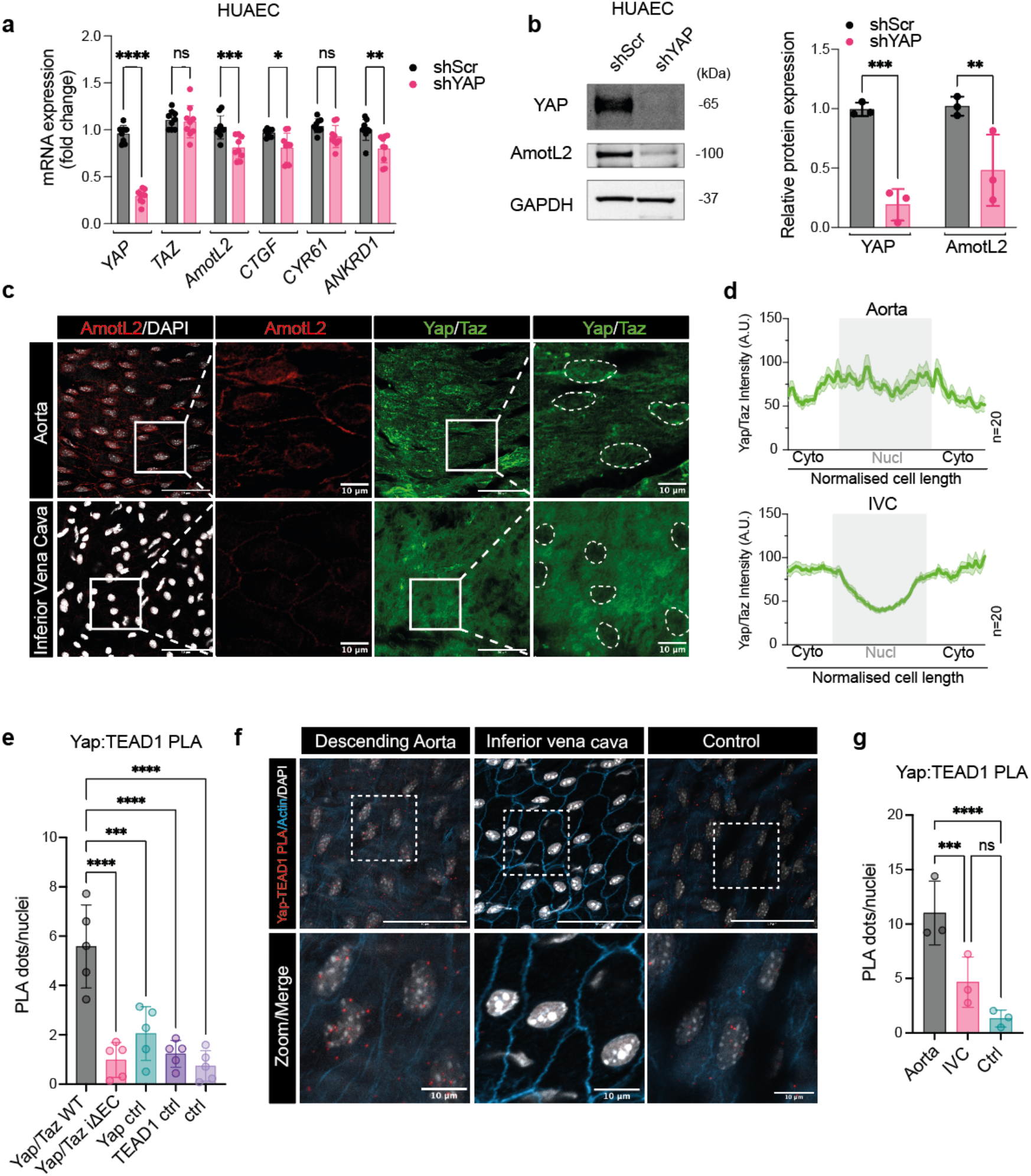
The murine aorta exhibits high AmotL2 expression and YAP activity compared to the vena cava. **a**, AmotL2 is transcriptionally regulated by YAP. Fold change in mRNA expression of *AmotL2* and known YAP target genes (*CTGF, CYR61* and *ANKRD1*) targets in HUAEC cells transduced with shScr or shYAP lentiviral vectors, analysed by qPCR and normalised to GAPDH. *n*=3 independent experiments, each with 3 technical replicates. (mean± s.d., 2way ANOVA with Dunnett’s multiple comparisons) **b**, Representative western blot showing YAP and AmotL2 expression in HUAEC cells transduced with shScr or shYAP lentivirus for 96h prior to immunoblot analysis. GAPDH was used as a loading control and normalisation for respective quantification shown in the right-hand panel, *n*=3 independent experiments. (mean± s.d., 2way ANOVA with Dunnett’s multiple comparisons) **c**, Confocal imaging of *en face* whole mount staining of the descending aorta (upper panel) and inferior vena cava (lower panel). AmotL2 is shown in red and YAP/TAZ in green. Nuclei are outlined by white dotted lines for clarity. Scale bar, 50µm. In zoomed panels, Scale bar, 10µm. **d**, Quantification of nuclear and cytoplasmic staining of YAP/TAZ. Histogram graph depicts the average intensity (mean± s.e.m.) of *n*=20 cells from one representative animal. A total of 60 cells per aorta or IVC were measured from *n*=3 mice. See supplementary figure 3b for full quantification. **e**, Quantification *ex vivo-en face* PLA of YAP-TEAD1 PLA of WT and YAP/TAZ iΔEC and indicated controls (images shown in full in SuppFig2c), where each data point represents an individual animal, (*n*=5 mice per group, mean± s.d., 2way ANOVA with Dunnett’s multiple comparisons) **f**, Representative images of *ex vivo-en face* PLA of YAP-TEAD1 interactions in the endothelium of the descending aorta and inferior vena cava. Positive interactions are indicated by red dots, nuclei in grey and actin in blue. **g**, Quantification of d, from *n=3* independent WT mice (mean± s.d., 2way ANOVA with Dunnett’s multiple comparisons). Scale bar, 50µm and 10µm, for 40x and indicated zoomed regions respectively.

### AmotL2 and YAP are highly expressed in the aorta compared to the vena cava

Previous work from our lab has shown that AmotL2 is highly expressed in aorta compared to inferior vena cava (IVC)^5^. As YAP transcriptionally regulates AmotL2, we hypothesised that this pattern of vascular expression may be due to increased YAP activity exhibited in the aorta^10,11,12^. *In silico* screening of the ENCODE RNA Annotation and Mapping of Promotors for Analysis of Gene Expression (RAMPAGE)^13,14^ indicated that the transcriptional activity of the promotor of YAP was most active in the thoracic aorta (SuppFig3a). We used immunofluorescence to visualise YAP and AmotL2 expression and localisation in the descending aorta and compared this to the IVC. Results indicated increased AmotL2 expression levels and greater nuclear YAP localisation in the descending aorta compared to the vena cava (Fig1c-d and SuppFig3b). To explore the increased nuclear YAP localisation of the aorta in more detail, we made use of the proximity ligation assay (PLA), to probe for the interaction between YAP and the DNA binding transcription factor, TEAD1. The proximity of both these proteins, is indicative of active nuclear YAP, which is able to indirectly bind to DNA via TEAD1 and drive transcription^11,15,16,17^. To validate the specificity of YAP-TEAD1 PLA on *ex vivo* aortic tissue, we utilised inducible endothelial specific Cdh5(BAC)^CreERT2^ transgenic mice crossed with *Wwtr1 flox/flox*; *Yap flox/flox* (herein referred to as Yap/Taz iΔEC). The genetic deletion of one of the targets of the PLA reaction (in this case Yap) resulted in the ablation of PLA signal in *ex vivo* samples from the aorta of Yap/Taz iΔEC mice (SuppFig3c). Quantification of YAP-TEAD1 PLA signal in Yap/Taz iΔEC mice, compared to their wild type (Yap/Taz WT) counterparts, indicated a significant reduction in PLA signal, comparable to single antibody, and double negative control conditions (Fig1e), suggesting high specificity of the technique. We next compared the aorta and vena cava for YAP-TEAD1 interactions using PLA as a readout of YAP activity. Results indicated increased PLA signal in the descending aorta, compared to the vena cava (Fig1f-g), confirming immunofluorescent stainings showing increased nuclear YAP in the aorta. These results suggest that the aorta exhibits increased YAP activity and consequently increased AmotL2 expression, compared to the inferior vena cava.

### Endothelial AmotL2 is regulated by YAP *in vivo*

To test whether Yap transcriptionally controls AmotL2 *in vivo*, and whether high AmotL2 expression of the aorta is Yap dependent, we performed whole mount immunofluorescent staining of AmotL2 and Yap on the aortae of WT and Yap/Taz iΔEC mice. Immunofluorescent staining showed a reduction in Yap/Taz intensity and a concomitant reduction in AmotL2 expression in Yap/Taz iΔEC mice (SuppFig3a and Fig2a). Within regions of the Yap/Taz iΔEC aorta we observed areas of endothelium which stained positively for Yap/Taz which were accompanied by higher expression levels of AmotL2, serving as an internal control within the Yap/Taz iΔEC aorta (Fig2b). Quantification indicated that this was consistent across the cohort of Yap/Taz iΔEC mice (Fig2b). These results confirm that Yap/Taz regulates endothelial AmotL2 expression within the aortic endothelium. Endothelial specific deletion of AmotL2 results in the depletion of junctional actin fibres of the aorta, leading to perturbation in endothelial nuclear morphology and abnormal endothelial sensing of shear flow^5^. Therefore, we hypothesised that as an upstream regulator of AmotL2, the deletion Yap/Taz would phenocopy the morphological dysfunction observed in AmotL2 iΔEC. Staining of the actin cytoskeleton and nucleus in the descending aorta indicated a loss of radial actin fibres in Yap/Taz iΔEC compared to WT Yap/Taz aortae (Fig2c) and more rounded nuclei, compared to the elongated morphology observed in Yap/Taz WT aorta (Fig2d). These results mirror *in vivo* endothelial cell morphology of AmotL2 iΔEC and suggest that Yap is required for AmotL2-dependent response to shear flow in the aorta.

**Fig 2.**
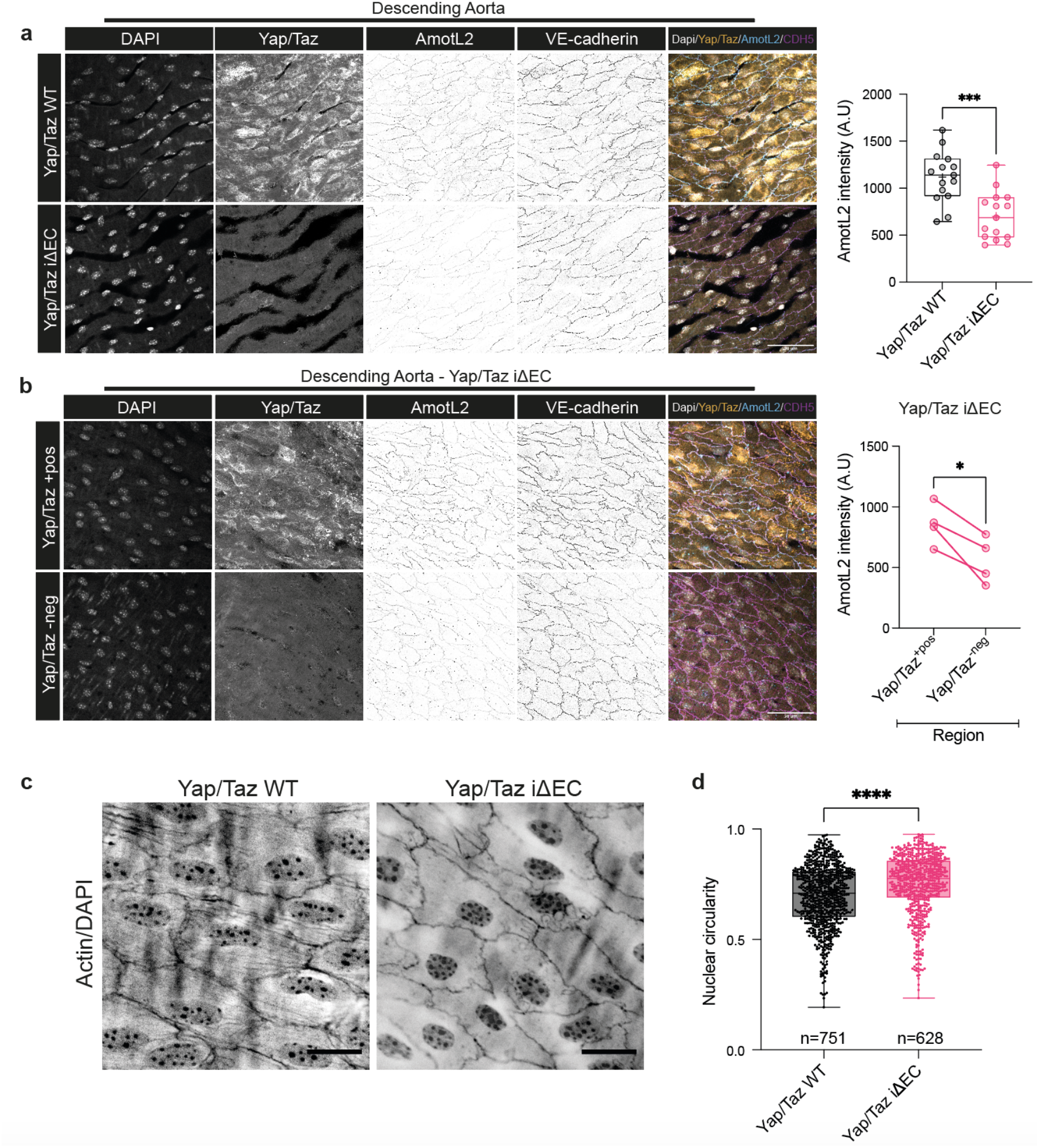
Deletion of endothelial Yap/Taz decreases AmotL2 expression and disrupts nuclear shape in aortic endothelial cells *in vivo*. **a**, Representative images of *en face* staining of AmotL2 in the descending aorta of both Yap/Taz WT and Yap/Taz iΔEC mice, *n*=5 per group. Nucleus (grey), Yap/Taz (yellow), Ve-cadherin (magenta). Scale bar, 50µm. Bar graph indicates quantification of AmotL2 fluorescent intensity, where each data point represents intensity profile from one image, 3 images/aorta (*n*=5 mice/group) were analysed, mean± s.d., Mann-Whitney). **b**, Representative images of regions of positive (+pos) and negative (-neg) Yap/Taz staining in Yap/Taz iΔEC mice and corresponding AmotL2 (blue), Ve-cadherin (magenta), and nuclear (grey) staining. Scale bar, 50µm. Graph indicates quantification of AmotL2 fluorescent intensity, where each data point represents average intensity profile from one mouse, (*n*=4 mice/group)(paired t-test). **c**, Representative images of *en face* staining of actin and the nucleus in endothelium of the descending aorta of both Yap/Taz iΔEC and Yap/Taz WT mice, *n*=5. **d**, Quantification shown in box and whisker plot indicates nuclear circularity, measured using ImageJ as described in the methods link. (*n=751* and *n=628* nuclei from 5 mice/group, mean± s.d., Mann-Whitney).

### AmotL2 regulates *YAP/TAZ* transcription

Previous work has suggested that AmotL2 negatively regulates YAP activity through modulation of the actin cytoskeleton^18^, upstream kinases of the Hippo pathway^19,20,^ and direct binding to YAP itself^20,21,22^. We therefore investigated AmotL2 mediated regulation of YAP in human endothelial cells. Intriguingly, the depletion of AmotL2 in HUVEC and HUAEC by lentiviral shRNA, resulted in a significant reduction in YAP protein levels when analysed by western blotting (Fig3a and Fig3b). Depletion of AmotL2 with two additional shRNA constructs with different target sequences showed similar results in HUVEC, with reduced YAP expression (SuppFig5a-b). In HUAEC, downregulation of YAP was only noted with one of the additional constructs (SuppFig5a-b). To test whether AmotL2 depletion specifically downregulated YAP, we performed western blotting using an antibody which detects both YAP and it’s paralog transcriptional coactivator, TAZ. Here the depletion of AmotL2 led to a significant ablation of YAP protein, however no observable effect was noted in TAZ protein levels (SuppFig5c), which was mirrored in the quantification (SuppFig5d). To investigate whether changes in YAP protein were attributable to transcriptional activity or protein stability, RT-qPCR was performed in AmotL2 depleted HUVEC and HUAEC. Depletion of AmotL2 led to a significant reduction in *YAP* mRNA in both endothelial cell lines, suggesting that *YAP* transcription is AmotL2 dependent (Fig3c and 3d). To further investigate the role of endothelial AmotL2 in regulating YAP activity, we performed western blotting for phosphorylated YAP and TAZ in AmotL2 depleted HUVEC. As before, a reduction in total YAP levels was observed in AmotL2 depleted HUVEC (Fig3e). No observable difference was noted in ser127 phosphorylation, however, when normalising and considering the depletion in total YAP, ser127 phosphorylation was found to increase significantly, suggesting decreased YAP activity (Fig3f). No observable difference was noted in TAZ ser89 phosphorylation upon depletion of AmotL2 (SuppFig5e). Additionally, subcellular fractionation and analysis of YAP localisation did not indicate any substantial difference in the nuclear:cytoplasmic ratio of YAP (Fig3g and quantified in SuppFig5f). To further validate AmotL2-dependent YAP-activity we performed ChIP-qPCR in AmotL2-depleted HUVEC and immunoprecipitated chromatin using a YAP-specific antibody. Analysis of the *CTGF* and *AmotL2* promotors by ChIP-qPCR indicated a reduction in YAP binding upon AmotL2 depletion, suggesting that endothelial YAP activity is reduced in the absence of AmotL2 (Fig3h, SuppFig5h-i). Collectively, these results suggest that transcription of *YAP* is AmotL2 dependent in endothelial cells, and that this impacts on YAP’s activity as a transcriptional co-activator.

**Fig 3.**
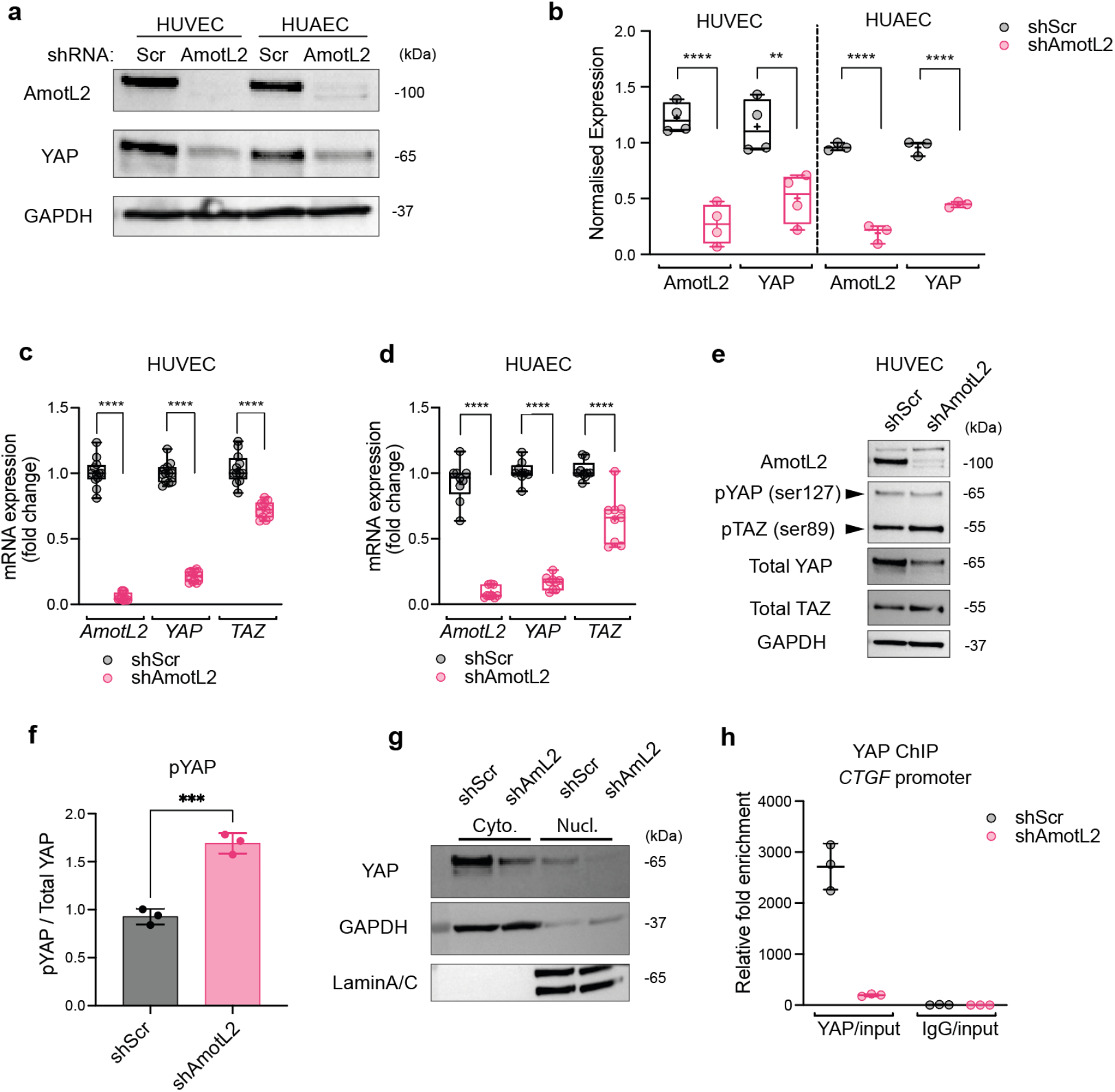
AmotL2 regulates *YAP/TAZ* transcription. **a**, Western blot analysis of AmotL2 and YAP in HUVEC and HUAEC cells 96h post-treatment with shScr or shAmotL2 lentivirus. GAPDH was used as a loading control. **b**, Quantification of AmotL2 and YAP protein levels, relative to GAPDH loading control. HUVEC *n*=4, HUAEC *n*=3 independent experiments, mean± s.d., 2way ANOVA with Dunnett’s multiple comparisons. **c**, SYBR green RT-qPCR of *AmotL2, YAP*, and *TAZ* relative to housekeeping gene *GAPDH*, in AmotL2 knockdown HUVEC and HUAEC cells. (*n*=4 independent experiments for HUVEC and *n*=3 for HUAEC, mean± s.d., 2way ANOVA with Dunnett’s multiple comparisons). **e**, Western blot analysis of indicated proteins HUVEC cells 96h post-treatment with shScr or shAmotL2 lentivirus. GAPDH was used as a loading control. **f**, Quantification of pYAP ser127 levels, relative to total YAP. (*n*=3 independent experiments, mean± s.d., Mann-Whitney). **g**, Representative western blot of nuclear:cytoplasmic fractionation detection of indicated proteins in HUVEC cells 96h post-treatment with shScr or shAmotL2 lentivirus. GAPDH and LaminA/C were used as positive and negative controls. **h**, ChIP showing YAP binding to *AmotL2* and *CTGF* promotor of shScr or shAmotL2 treated HUVEC. ChIP qPCR was performed using SBYR green reagents and quantification was normalised to an IgG control antibody. Plot shown is a representative experiment from *n=3* independent experiments (Supplemental figure 5h shows two further independent experiments). Each data point represents a technical repeat within one independent experiment (performed in triplicate) (mean± s.d.).

### AmotL2 regulates Yap expression of the aortic endothelium

To test whether the Yap expression was also AmotL2 dependent *in vivo*, we utilised Cdh5(PAC)^CreERT2^ transgenic mice^23^ crossed with *amotl2*^flox/flox^ mice and ROSA26-EYFP reporter mice to specifically delete endothelial AmotL2 in adult mice (herein referred to as AmotL2 iΔEC). As AmotL2 expression and Yap activity was found to be expressed in the aorta (Fig1), we tested whether the deletion of AmotL2 in the endothelium of the aorta resulted in the downregulation of Yap in this context. Using VE-cadherin to confirm the EC identity, immunofluorescence indicated complete ablation of Yap expression in a number of cells of the AmotL2 iΔEC aorta (Fig4a). Using the EYFP reporter for AmotL2 deletion allowed the identification of KO cells upon recombination, which indicated EYFP positive cells with reduced Yap/Taz expression, compared to neighbouring EYFP negative cells, exhibiting WT Yap/Taz expression levels (Fig4b, quantification in Fig4c showing reduced Yap/Taz (green histogram) expression in EYFP positive cells (purple histogram)). Importantly, staining of Yap/Taz in the descending aorta of tamoxifen injected CreERT2 EYFP positive mice indicated homogenous levels of Yap/Taz expression across both EYFP positive and negative cells (SuppFig6c). The inferior vena cava of AmotL2 iΔEC animals did not indicate a reduction in Yap/Taz staining (SuppFig6a). As Yap/Taz appeared cytoplasmic and largely inactive in the IVC, and the reported role of AmotL2 as a negative regulator of Yap/Taz activity, we analysed Yap/Taz nuclear:cytoplasmic ratio of AmotL2 iΔEC, which indicated no change in total levels, or localisation of Yap/Taz compared to AmotL2 WT IVC (SuppFig6b). Single cell sequencing of murine lungs highlights the ubiquitous and varying expression of *AmotL2* in tissues^24^,^25^ (SuppFig6d-e). Western blotting of total lung lysates from AmotL2 WT and AmotL2 iΔEC mice indicated a significant reduction in AmotL2 and Yap protein levels, despite the heterogenous nature of the whole lung lysates (Fig4d, quantification Fig4e-f). These results show that deletion of endothelial AmotL2 results in the downregulation of Yap *in vivo*, mirroring *in vitro* findings. To further investigate the relationship between *AmotL2* and *YAP* expression in human vascular samples, we explored transcript levels in human aortic samples surgically resected from patients undergoing reconstructive surgery of the abdominal aorta. Transcriptomic array data of the medial layer of resected aorta with intact endothelium showed a correlation between *YAP* and *AmotL2* (Fig4g). Correlation of *TAZ* mRNA and *AmotL2* was however, not significant (Fig4h) indicating the *YAP* specific coregulation with *AmotL2*. Overall, these data indicate that endothelial YAP expression *in vivo* is AmotL2 dependent, suggesting a positive feedback loop of regulation.

**Fig 4.**
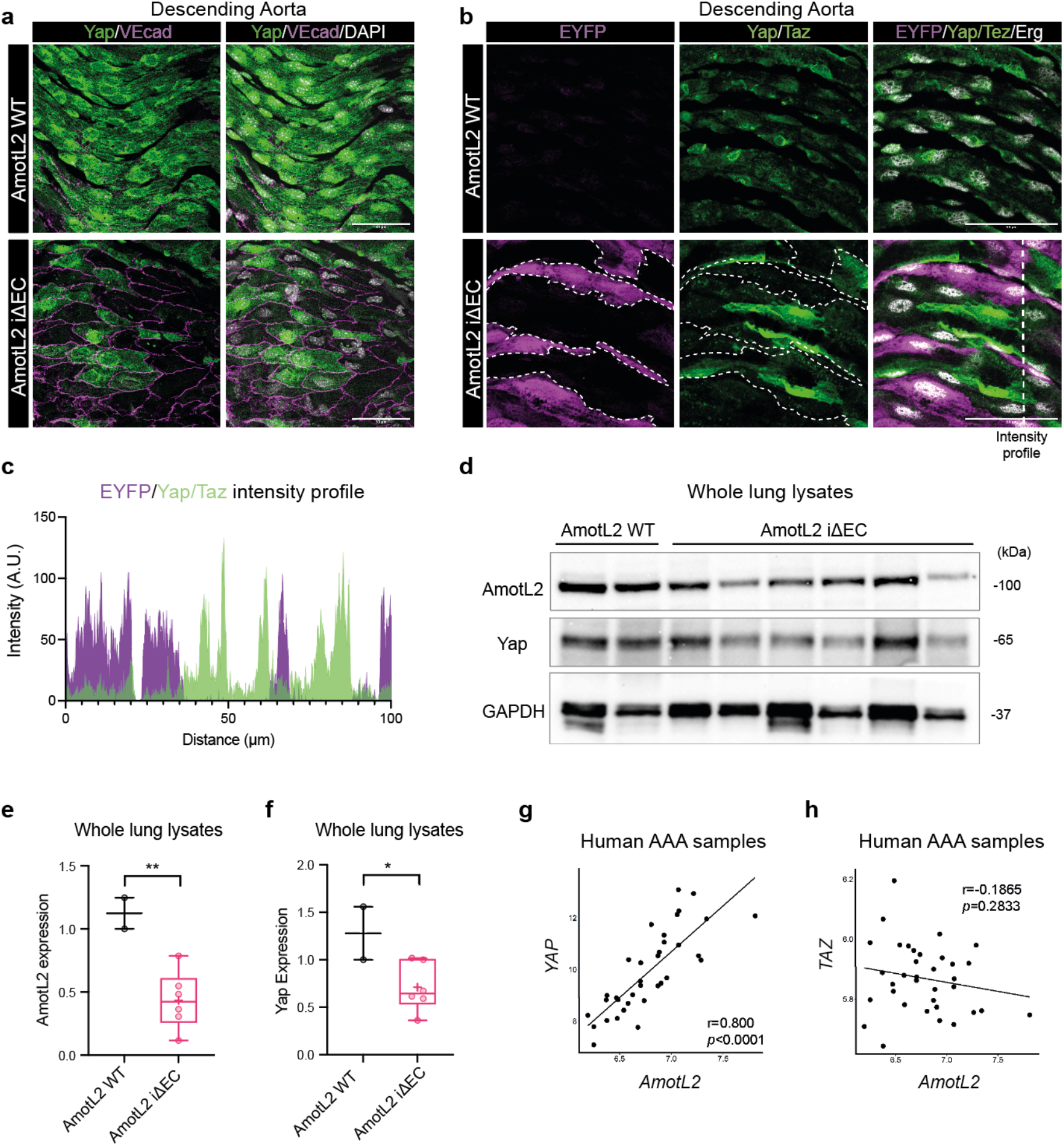
AmotL2 regulates aortic endothelial YAP *in vivo*. **a**, Representative images of *en face* staining of Yap (green) and VE-cadherin (magenta) in the descending aorta of both AmotL2 WT and AmotL2 iΔEC mice. Nucleus (grey), Scale bar, 50µm. Images are representative of *n=3* mice/group. **b**, Representative images of *en face* staining of EYFP (magenta), Yap/Taz (green), and Erg (grey) in the descending aorta of both AmotL2 WT and AmotL2 iΔEC mice. EYFP positive cells are outlined for clarity. Scale bar, 50µm. Images are representative of *n=3* mice/group. **c**, Line profile (indicated by dashed white line in the bottom right-hand panel of b) of immunofluorescent intensity of EYFP and Yap/Taz of AmotL2 iΔEC aorta. **d**, Western blot analysis of AmotL2, Yap protein expression from AmotL2 WT (*n*=2) and AmotL2 iΔEC (*n*=6) whole lung lysates. GAPDH was used as a loading control. AmotL2 WT (*n*=2) and AmotL2 iΔEC (*n*=6). **e**, Box plot indicates quantification of AmotL2 expression relative to GAPDH loading control. (AmotL2 WT (*n*=2) and AmotL2 iΔEC (*n*=6), mean± s.d., unpaired t-test). **f**, As in e for Yap protein expression. **g**, *AmotL2* correlates with *YAP* in human medial aortic tissue. The correlation between *AmotL2* mRNA and *YAP* and *TAZ* (**h**) mRNA of aneurysms in AAA patients (n=35). *AmotL2* expression level was based on the expression of the first exon from 3’ end, detected by specific exon probe, which represents the full-length isoform of *AmotL2*.

### AmotL2 regulates YAP-dependent proliferation

YAP is a key regulator of cellular proliferation^26,27,28,29^. To examine the functional implication of AmotL2-dependent YAP regulation, we tested whether AmotL2 depletion impacted on EC proliferation. ECs were subjected to AmotL2 depletion and live cell imaging in order to analyse cell growth over time. Growth curves of shScr and shAmotL2 treated HUVEC indicated a significant reduction in confluency over a 68h time course (Fig5a), in line with previously observed findings^30^. The proliferation of untransformed cells is limited by negative feedback through contact inhibition, which in part is controlled by YAP localisation^31^. As such, analysis of proliferation by EdU incorporation of subconfluent and confluent HUVEC endothelial cells indicated a greater percentage of cells in S-phase in subconfluent compared to confluent conditions (Fig5b, SuppFig7a). Depletion of AmotL2 in HUVEC significantly reduced the percentage of cells in S-phase under subconfluent conditions when plated on plastic (Fig5b), results which were mirrored on 50kpa hydrogels (SuppFig7b), albeit reduced in comparison to the stiffer plastic conditions shown in Fig5b. Importantly, treatment of HUVEC with the YAP-TEAD competitive inhibitor, Verteporfin (VP), mimicked results of *in vitro* AmotL2 depletion, both in growth kinetics analysed by live cell imaging (Fig5c), and EdU incorporation (SuppFig7c and Fig5d), indicating the reliance of EC proliferation on YAP-TEAD interactions in driving proliferation.

**Fig 5.**
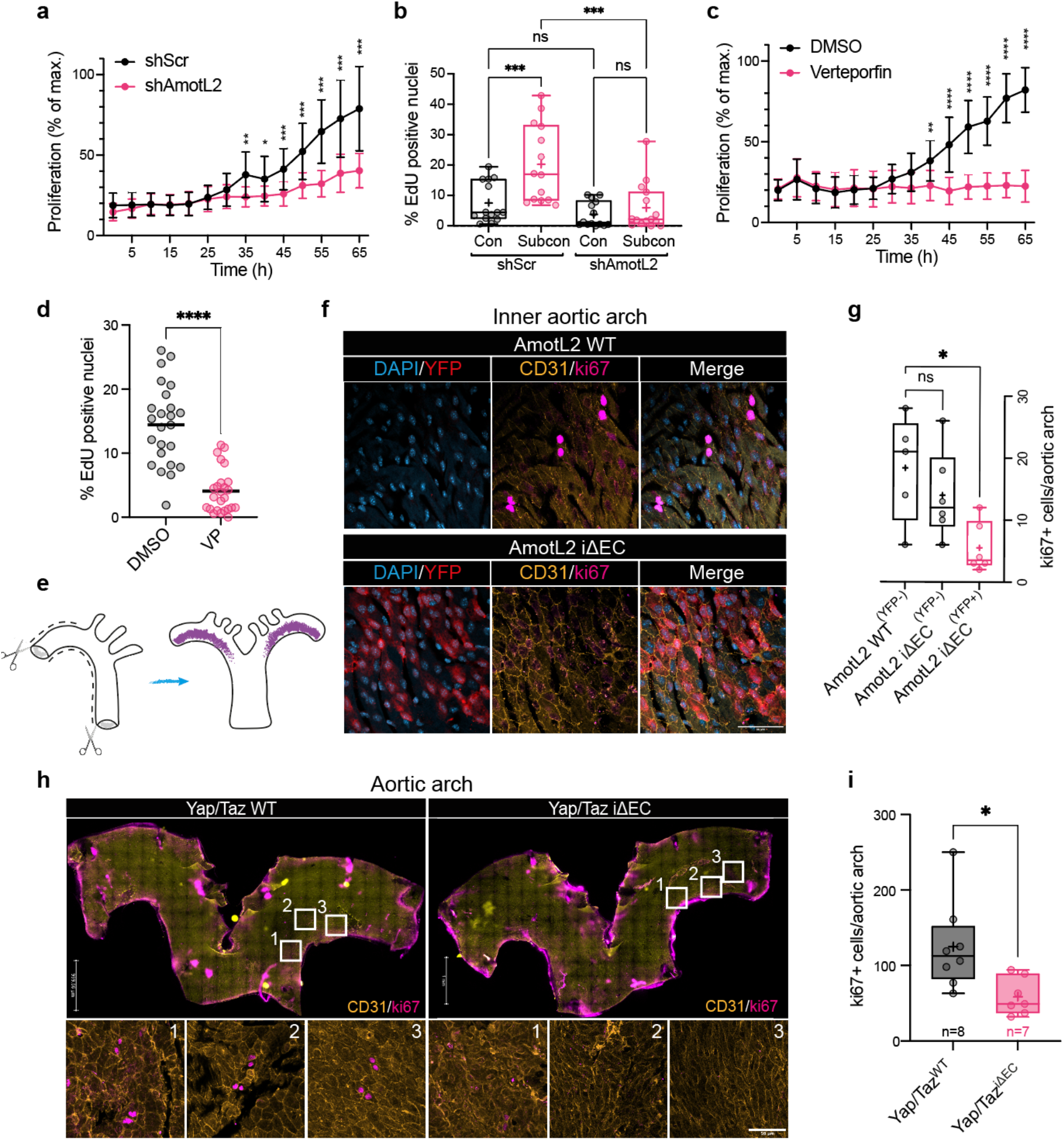
Endothelial proliferation is regulated by AmotL2 and YAP. **a**, Proliferation of HUVEC replated to gelatin coated plastic following 48h post-lentiviral transduction with shScr or shAmotL2 lentivirus. *n*=3 independent experiments (mean± s.d., 2way ANOVA with Sidak’s multiple comparisons) error bars indicate standard deviation. **b**, Box plots showing quantification of EdU, where % of EdU positive cells was calculated against total number of cells stained with Hoechst. Each data point represents one field of view from *n*=3 independent experiments. (mean± s.d., 2way ANOVA with Dunnet’s multiple comparisons). **c**, Proliferation of HUVEC plated to gelatin coated plastic following and treated with 0.2ug/ml Verteporfin or DMSO vehicle *n*=3, error bars indicate standard deviation. **d**, Quantification of EdU positive HUVEC treated with 0.2ug/ml Verteporfin or DMSO vehicle (48h). Cells were counterstained with Hoechst and quantified where % of EdU positive cells was calculated against total number of cells stained with Hoechst. (Each data point represents one field of view from *n*=4 independent experiments, bar indicates mean, Mann-Whitney) **e**, Schematic depicting dissection of whole aorta for en face whole mount staining. Purple region indicates the inner aortic arch. Schematic drawn using Adobe Illustrator (v25.4.1). **f**, Representative images of the inner aortic arch of AmotL2 WT and AmotL2 iΔEC, *en face* whole mount stained for YFP (red), nucleas (blue), CD31 (yellow), ki67 (magenta), using ChrisLUT BOP palette (https://github.com/cleterrier/ChrisLUTs). Scale bar, 50µm. Images representative of *n=5* mice/group. **g**, Quantification of total ki67 and CD31 positive cells per aortic inner arch from AmotL2 WT (YFP −neg) *n*=5, AmotL2 iΔEC (YFP −neg) *n*=5, (YFP +pos) *n*=5. (mean± s.d., 2way ANOVA with Dunnet’s multiple comparisons). **h**, Representative images of the inner aortic arch of Yap/Taz WT and Yap/Taz iΔEC, *en face* whole mount stained for CD31 (yellow), nucleus (blue), and ki67 (magenta) Scale bar, 50µm.**i**, Quantification of total ki67 and CD31 positive cells per aortic inner arch from Yap/Taz WT, (*n*=5), Yap/Taz iΔEC, (*n*=5). (mean± s.d., Mann-Whitney).

To investigate whether AmotL2 regulates proliferation *in vivo*, we focused on a subpopulation of proliferating ECs present within the inner arch of the adult aorta^32,33^ using *en face* whole mount staining (Fig5e). Staining of the inner aortic arch of AmotL2 WT mice with the marker of proliferation, ki67, the EC specific marker CD31 and EYFP indicated the presence of ki67 positive ECs (Fig5f). The induction of AmotL2 deletion, indicated by the expression of the EYFP reporter, was accompanied by a reduction in ki67 positive ECs in the inner aortic arch (Fig5f, lower panel). Quantification indicated a significant reduction in ki67+/EYFP+ ECs in AmotL2 iΔEC compared to AmotL2 WT, EYFP-littermates (Fig5g). To confirm that the proliferation of the ECs of the inner aortic arch is indeed dependent on YAP signalling we stained the inner aortic arches of WT Yap/Taz mice and Yap/Taz iΔEC with ki67 and CD31 and analysed ki67+/CD31+ cells. Staining indicated a reduction in proliferating ECs of Yap/Taz iΔEC arches, compared to Yap/Taz WT aortae (Fig5h) which was confirmed by quantification of ki67+/CD31+ cells (Fig5i). Collectively, these results indicate that mechanically induced endothelial proliferation is regulated by a AmotL2-YAP axis, the absence of which leads to suppressed proliferation both *in vitro* and *in vivo*.

### AmotL2 regulates *YAP* transcription by histone methylation

To further understand how AmotL2 regulates *YAP* transcription, we focused on investigating changes to epigenetic markers associated with transcriptional repression. In the initial reporting of the cloning of YAP, Sudol et al showed that *YAP* was not expressed in human PBMCs^34^, nor in the murine spleen^35^. Using the human protein atlas, we confirmed that the expression of *YAP* in blood cells was virtually undetectable across several organs analysed by scRNAseq. SuppFig8a depicts *YAP* expression from one example of human scRNAseq of breast tissue showing reduced expression of *YAP* in the blood and immune cells compared to other cell types within the dataset. Using a single-cell atlas of chromatin accessibility in humans (http://catlas.org/catlas_hub/), we confirmed that chromatin around the promotor of *YAP* exhibited a closed-condensed conformation in CD4+, CD8+, and naïve T cells and NKT cells, compared to the open chromatin exhibited by ECs of a number of organs (SuppFig8b). Open chromatin allows binding of RNA polymerase II (Pol II) to actively transcribe mRNA from the accessible DNA template. We therefore tested whether the downregulation of *YAP* transcription was due to differential binding of Pol II to the *YAP* promotor. Immunoprecipitation of chromatin from AmotL2 depleted HUVEC using a Pol II specific antibody indicated a significant reduction in binding to the *YAP* promotor, compared to shScr and IgG controls (Fig6a). DNA methylation is widely accepted as a marker of transcriptional repression. Recently, differential *YAP* promotor methylation of cells cultured on soft vs stiff matrices was reported^36^. Further analysis of published data indicated increased methylation of the *YAP* promotor in immune cells compared to other *YAP* expressing cell types (Depmap.org) (SuppFig8c).

**Fig 6.**
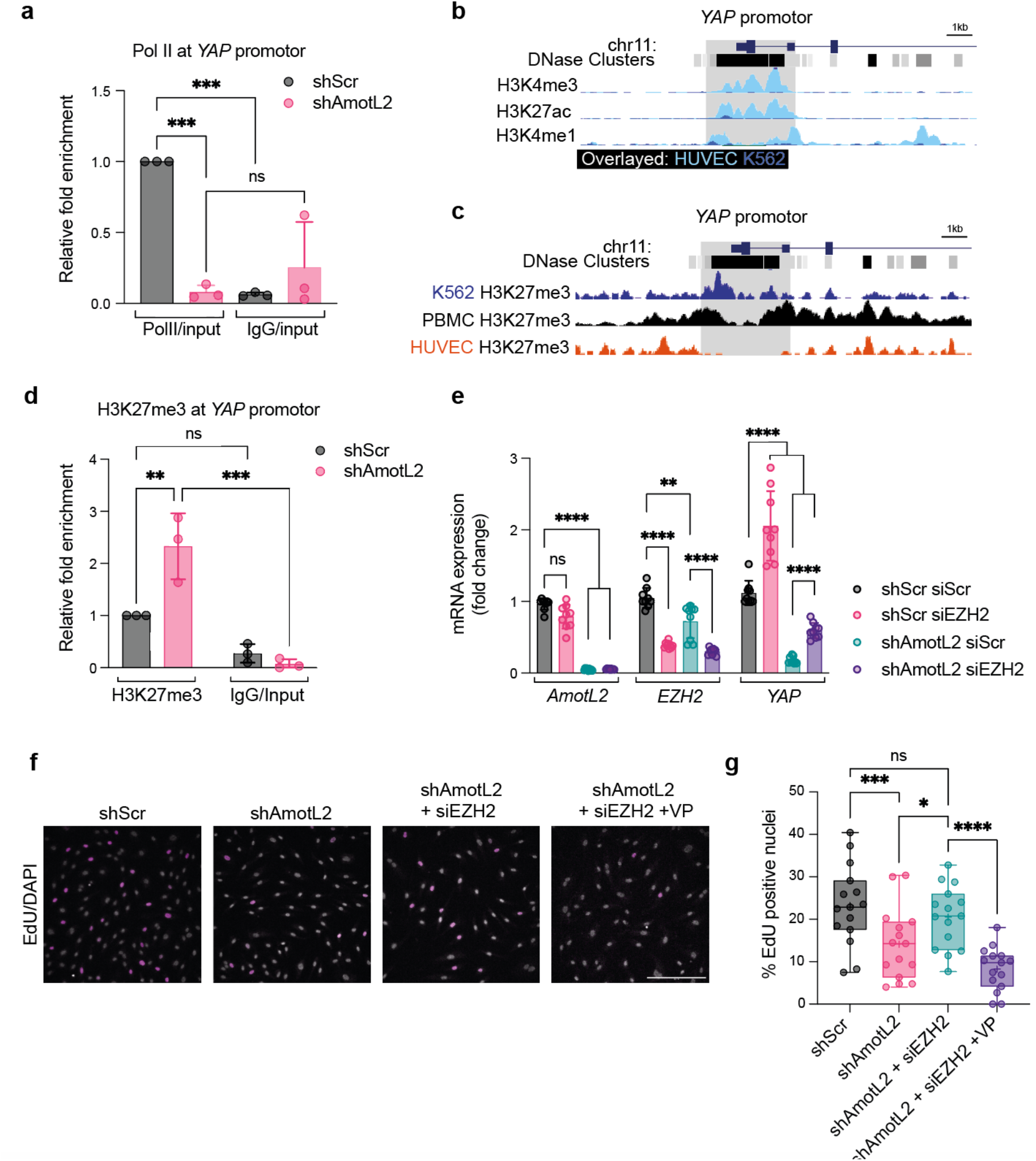
AmotL2 regulates H3K27me3 to modulate *YAP* promotor activity. **a**, ChIP showing Pol II binding to the *YAP* promotor of shScr or shAmotL2 treated HUVEC. ChIP qPCR was performed using SBYR green reagents and quantification was normalised to an IgG control antibody. (*n=3* independent experiments, mean± s.d., 2way ANOVA with multiple comparisons). **b**, Screen grab of the UCSC browser displaying genomic tracks showing published ChIP-seq data of H3K4me3, H3K27ac and H3K4me1 of ChIP-seq data across HUVEC and K562 (overlayed) within the *YAP* promotor (Data sources are referenced in the methods). **c**, Screen grab of the UCSD browser displaying genomic tracks showing ChIP-seq data of H3K27me3 of ChIP-seq data across K562, PBMC and HUVEC within the *YAP* promotor (Data sources are referenced in the methods). **d**, ChIP-qPCR analysis of H3K27me3 binding to the *YAP* promotor of shScr or shAmotL2 treated HUVEC. ChIP qPCR was performed using SBYR green reagents and quantification was normalised to an IgG control antibody. Plot shown is representative of *n=3* independent experiments, mean± s.d., 2way ANOVA with Šídák’s multiple comparisons. Each data point represents an independent experiment. **e**, Codepletion of EZH2 and AmotL2 rescues *YAP* mRNA expression. RT-qPCR of indicated targets in HUVEC cells where knockdown of EZH2 using siRNA followed by shRNA knockdown using shScr or shAmotL2. (*n=3* independent experiments, mean± s.d., 2way ANOVA with Tukey’s multiple comparisons). **f**, Representative images of shScr, shAmotL2, shAmotL2 + siEZH2, shAmotL2 + siEZH2 + Verteporfin (VP) at 0.2ug/ml treated HUVEC 72h post infection with lentiviral vectors, 72 h post siRNA transfection, replated to gelatin coated plastic in subconfluent conditions. Incorporated EdU was detected with secondary antibodies and counterstained with Hoechst. Scale bar, 250µm. **g**, Quantification of (f), where each data point represents one field of view (5 taken per condition) from *n*=3 independent experiments, mean± s.d., 2way ANOVA with Tukey’s multiple comparisons).

However, global analysis of methylation of AmotL2 depleted HUAEC cells did not indicate global differential methylation and so we turned our attention to histone modifications (SuppFig8d).

A plethora of epigenetic modifications to histones are responsible for transcriptional regulation of gene promotors, many of which are temporal and cell-type specific. We therefore used existing ChIP-seq data to compare known *YAP*-expressing and non-expressing cell types, to ascertain the main contributors to epigenetic regulation of *YAP*, and then investigate if AmotL2 modulated these. Analysis of the *YAP* promotor within UCSC genome browser and ENCODE database revealed enrichment for the markers of active euchromatin; H3K4me3, H3K27ac and H3K4me1 in HUVEC, compared to the low *YAP*-expressing leukemic cell line K562 (Fig6b). Further investigation indicated that the repressive histone marker H3K27me3, was enriched at the *YAP* promotor of PBMCs and K562 but was greatly reduced in HUVEC (Fig6c). We validated these datasets by exploring the pattern of histone modifications within the promotors of genes known to be exclusively expressed by ECs (*VEGFR2*)(SuppFig8e), immune cells (*CD45*)(SuppFig8f), or epithelial cells (*CDH1*)(SuppFig8g), which indicated expected patterns of enrichment for the active (H3K4me3, H3K27ac and H3K4me1) and repressive (H3K27me3) modifications.

We therefore hypothesised that AmotL2 regulates *YAP* transcription through modulation to H3K27 methylation. Depletion of AmotL2 and subsequent pulldown of chromatin using H3K27me3 antibody and analysis of the *YAP* promotor by qPCR revealed an enrichment of H3K27me3 compared to shScr conditions, indicating that repression of *YAP* may be due to increased repressive histone modifications in HUVEC (Fig6d) and HUAEC (SuppFig9). The catalytic subunit EZH2, forms part of the Polycomb Repressive Complex (PRC) and catalyses the methylation of H3K27^37,38,39,40^. We therefore performed a co-depletion of EZH2 and AmotL2 using siRNA, followed by lenti-shRNA respectively, to test whether we could prevent the repression of *YAP* transcription by EZH2 mediated methylation of H3K27. RT-qPCR results showed that co-depletion of AmotL2 and EZH2 led to a partial rescue in *YAP* mRNA (Fig6e), compared to lenti-shScr and Scr-siRNA conditions, indicating that AmotL2 regulates EZH2 and subsequently H3K27 methylation to modulate *YAP* promotor activity. Importantly, the depletion of EZH2 was able to rescue the proliferative capacity of shAmotL2 HUVEC, an effect that was abrogated by treatment with VP, suggesting that the rescue in proliferation in EZH2-AmotL2 co-depleted HUVEC was indeed due to increased YAP activity (Fig6f, quantification Fig6g). Collectively, these results indicate that the junctional molecule AmotL2 regulates *YAP* transcription through histone modifications to H3K27 via EZH2 activity.

### Chromatin accessibility is regulated by AmotL2

The notion of nuclear shape as a determinant of chromatin landscape and transcriptional activity is becoming increasingly evident^41^. Deletion of EC AmotL2 leads to mis-shapen nuclei of aortic ECs both *in vitro* and *in vivo*^5^, and in agreement with these data, we observed increased presence of nuclear wrinkles in AmotL2 knockdown HUAEC stained for a major structural component of the inner nuclear membrane, lamin A/C (Fig7a). We therefore investigated whether AmotL2 is involved in the regulation of lamins in maintaining nuclear morphology using subcellular fractionation and analysis of lamin A/C by western blotting, which showed a reduction specifically in lamin A expression (Fig7b). However, no change in *LAMINB1* was noted (SuppFig10a). We then asked whether regulation of nuclear shape through maintenance of lamin A by AmotL2 impacts on chromatin accessibility. To do this, we performed genome-wide transposase-accessible chromatin sequencing (ATAC-seq) on shScr and shAmotL2 knockdown in HUVEC and HUAEC cells. Figure 7c-d indicates volcano plots of peak accessibility in HUVEC and HUAEC cells, highlighting the widespread changes to chromatin accessibility when comparing AmotL2 depleted cells to shScr controls. As expected, the distribution of peaks detected indicated the enrichment of peaks in gene promotors and distal intergenic regions (Fig7e-f). Despite overall differences in the pattern of AmotL2 regulated chromatin conformation between HUVEC and HUAEC, KEGG pathway analysis of transcriptional start sites (TSSs) in which peak accessibility was reduced showed several overlapping pathways (Fig7g-h, overlap between groups in bold). Notably, pathways in fluid shear stress and atherosclerosis, adherens junctions and regulation of the actin cytoskeleton appeared in the top 20 most significant less accessible pathways, in line with the proposed role of AmotL2 as a junctional sensor of shear stress. To directly investigate whether transcriptional regulation of *YAP* was due to changes in chromatin accessibility around the TSS of *YAP*, we examined shScr and shAmotL2 ATAC-seq overlayed with published ChIP-seq data of Pol II, H3K27ac, H3K4me3, EZH2 and H3K27me3, using the IGV gene browser. ATAC-seq data of shScr and shAmotL2 in both HUVEC and HUAEC showed a bimodal peak of accessibility, in which the downstream peak (highlighted in grey, Fig7i-j) aligned with Pol II binding, markers of active histones (H3K4me3 and H3K27ac), and reduced EZH2 and H3K27me3 repressive histone markers. While little difference was noted in the first peak of this bimodal chromatin accessibility in shScr compared to shAmotL2, a noticeable reduction in accessibility was observed in the second peak position (Fig7i-j). Analysis of ranked ‘less-accessible peaks’ indicated *YAP* as a TSS with significantly reduced accessibility in HUVEC (log_2_ fold change −1.1596, *P*=0.0174 (Supplementary table 1). Additionally, differential pathways associated with changes in peak accessibility in shScr vs shAmotL2, grouped by KEGG pathway analysis showed the ‘Hippo signalling pathway’ in HUVEC (SuppFig10b). To directly investigate whether the changes in chromatin in this specific area reduced binding of Pol II, we performed ChIP qPCR on the region of interest within the *YAP* promotor (SuppFig10c) using a Pol II antibody. Results showed that the knockdown of AmotL2 reduced Pol II binding to this specific region of the *YAP* promotor (SuppFig10d), suggesting that AmotL2-dependent epigenetic regulation of the *YAP* promotor may be due to genome-wide changes to chromatin landscape accessibility.

**Fig 7.**
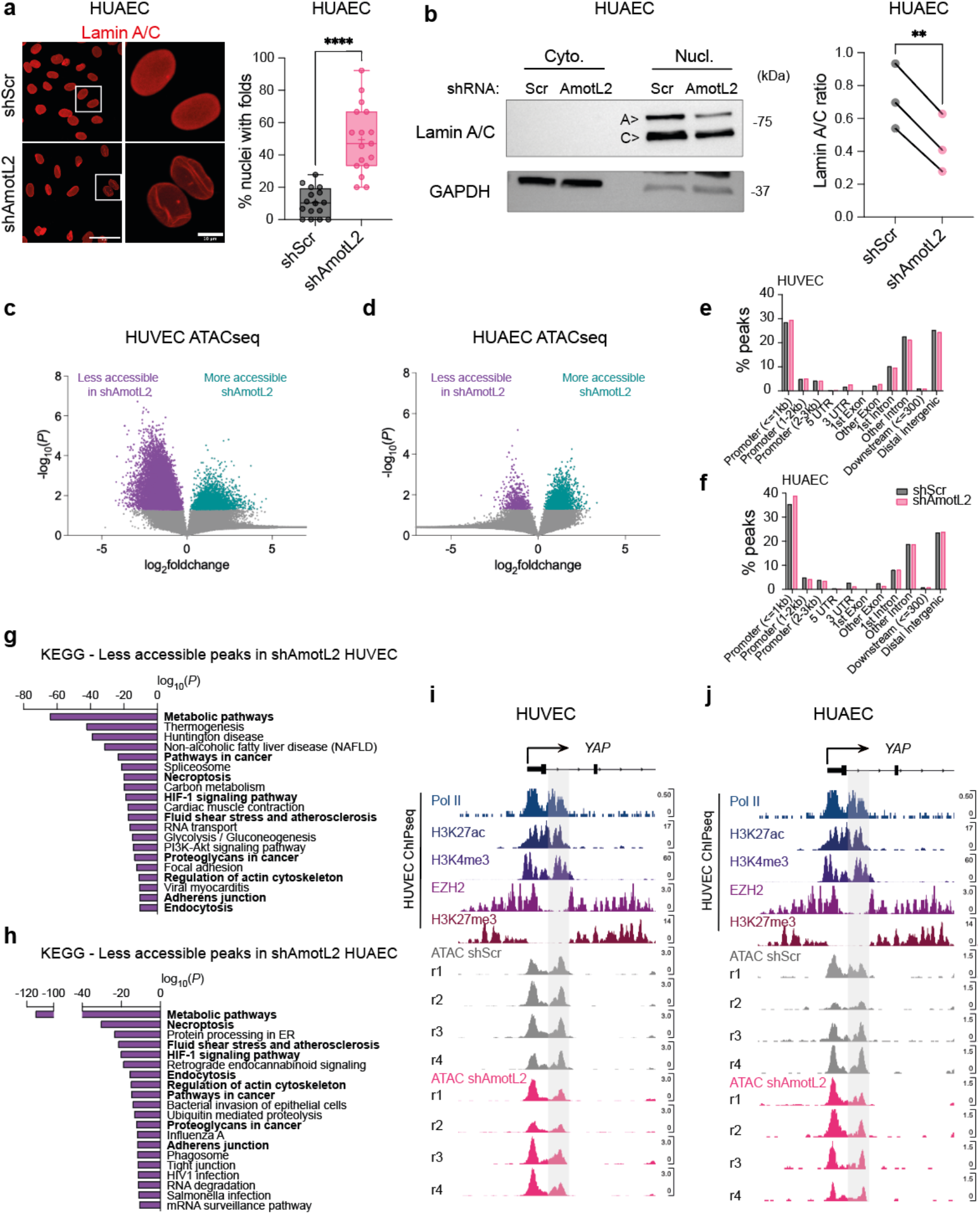
Chromatin accessibility is regulated by AmotL2. **a**, Immunofluorescent images showing lamin A/C in HUAEC with knockdown of AmotL2 or shScr control lentiviral vectors. Images are representative of *n*=3 independent experiments. Quantification indicates the number of nuclei with folds or wrinkles, calculated as a percentage of the total number of nuclei within a field of view. Each data point represents one field of view (5-7 taken per condition) from *n*=3 independent experiments, (mean± s.d., Mann-Whitney). **b**, Representative western blot of nuclear:cytoplasmic fractionation and detection of lamin A/C in HUAEC cells 96h post-treatment with shScr or shAmotL2 lentivirus. GAPDH were used as positive control for the cytoplasmic fraction. Quantification of lamin A/C ratio between matched shScr and shAmotL2 HUAEC cells samples. *n=3* independent experiments, paired t-test. Volcano plot indicating fold change in accessibility of HUVEC (**c**) or HUAEC (**d**) peaks of shScr control compared to shAmotL2. Peaks in purple show those that are less accessible in shAmotL2, and more accessible in shScr, while peaks shown in green are more accessible in shAmotL2 and less accessible in shScr. Grey points in the volcano plots represent peaks that are not significantly (*P*>0.05) changed between each group. **e-f**, Distribution of ATACseq peaks in different regions of the genome of both HUVEC and HUAEC. **g-h**, KEGG pathway analysis of peaks that are less accessible in HUVEC (**g**) and HUAEC (**h**) where genes that are involved in ‘Fluid shear stress and atherosclerosis’ and ‘actin regulation’ are less accessible in both cell lines, and so are highlighted in bold. **i-j**, IGV browser view of the *YAP* promotor showing publicly available ChIPseq data of Pol II, H3K27ac, H3K4me3, EZH2 and H3K27me3, alongside ATACseq data of HUVEC (**i**) and HUAEC (**j**) treated with shScr or shAmotL2. r indicates independent biological replicates, of which there are *n=*4 per condition.

## Discussion

Over the last decade much focus has been given to the localisation and activity of YAP and its downstream effect on transcription in response to mechanical stimuli. Previous work shows that AmotL2 is a negative regulator of YAP activity, through several mechanisms including direct binding and the modulation of upstream kinases of the Hippo pathway^18,19,20,21,22^. Our results show evidence that the depletion of AmotL2 leads to the downregulation of *YAP* mRNA, indicating a new model of YAP regulation by AmotL2, through modulation of transcription itself. Emerging evidence indicates the importance of transcriptional regulation of the Hippo pathway. Force dependent epigenetic regulation of the *YAP* promotor by DNA methylation occurs in gastric cancer cells cultured on soft vs. stiff matrices^36^ leading to reduced *YAP* mRNA. Whilst we could not detect changes to DNA methylation, we did observe alterations in histone modifications in the *YAP* promotor that were dependent on AmotL2. Kaukonen et al first described a role for JMJD1A, a histone demethylase, in a model of stiffness dependent regulation of *YAP/TAZ* transcription through modulation to H3K27ac^42^. More recently, changes in viscosity were associated with transcriptional changes to *YAP/TAZ* signalling^43^, but the precise mechanism of how this transcriptional regulation is elicited is unknown. These lines of evidence suggest that mechanical stimuli play a direct role in regulating *YAP* transcription. Our own results show that the junctional scaffold protein, AmotL2, plays a key role in the regulation of *YAP* transcription through regulation of nuclear shape, chromatin landscape, and H3K27me3 of the *YAP* promotor with functional consequence for endothelial cell proliferation.

AmotL2 was found to modulate EZH2 activity and downstream H3K27 methylation. H3K27me3 is mechanically sensitive^44^, and is regulated by fluid shear flow where disturbed flow of the inner aortic arch maintains high H3K27me3 and EZH2 expression^45^. Of interest, YAP has been shown to regulate recruitment of EZH2 to repress key regulators of the cell cycle and promote proliferation^46^. We therefore speculate that depletion of AmotL2, and a concurrent reduction in YAP may alleviate YAP-EZH2 dependent suppression of cell cycle regulators, leading to reduced proliferation. While we suggest that AmotL2 may alter EZH2 activity to mediate H3K27 methylation at the *YAP* promotor, the exact mechanism of how AmotL2 regulates EZH2 activity and the causal link to overall changes in chromatin, and the transcription of other genes remains to be determined.

Intriguingly, our findings show that AmotL2 regulates nuclear architecture and chromatin accessibility. This leads to further questions as to how a junctional molecule may influence the nucleus in such a way.

Previous work from our laboratory indicates that AmotL2 interacts with components of the nuclear membrane, namely lamin A, in ECs exposed to flow, suggesting an actin cytoskeleton derived link between cell-cell junctions and the nucleus^5^. A direct mechanical link between the nuclear membrane and integrins is well established, allowing the transduction of extracellular forces to the nucleus and subsequently shaping chromatin^41^. Additionally, recently pre-printed work has shown that disruption of the nuclear envelope LINC complex component SUN1 leads to dysregulation of endothelial cell junctions via the microtubule network, highlighting the existence of such a connection^47^. Whether the modulation of chromatin by AmotL2 is due to a direct connection from the cell-cell junctions and the nuclear membrane, or an indirect regulation of lamin A remains an open question.

YAP was also recently implicated in maintaining nuclear integrity through the actin cap by transcriptional regulation of *ACTR2* and the structural nuclear membrane protein *LAMINB1*^48^. In agreement with this phenotype, our findings indicated perturbed nuclei and a reduction in actin fibres in the aortic endothelium of Yap/Taz iΔEC, and phenocopy that of AmotL2 iΔEC which also display reduced radial actin fibres and nuclear dysmorphia^5^ and increased nuclear wrinkling *in vitro*. As AmotL2 depletion did not alter *LAMINB1* levels, we speculate that changes to nuclear morphology and chromatin are AmotL2-specific and are not a consequence of reduced YAP expression. However, we cannot rule out the downstream effects of chromatin regulation are partly a consequence of dysregulation of YAP’s ability to regulate chromatin accessibility through SWI/SNF and BRD4 interactions^49,50^.

Overall, our current model indicates that AmotL2 expression is critical for nuclear morphology, chromatin accessibility, and histone modifications within the *YAP* promotor (Fig8). To our knowledge, this represents the first example of a cell junctional molecule which influences epigenetic regulation of *YAP*.

**Fig 8.**
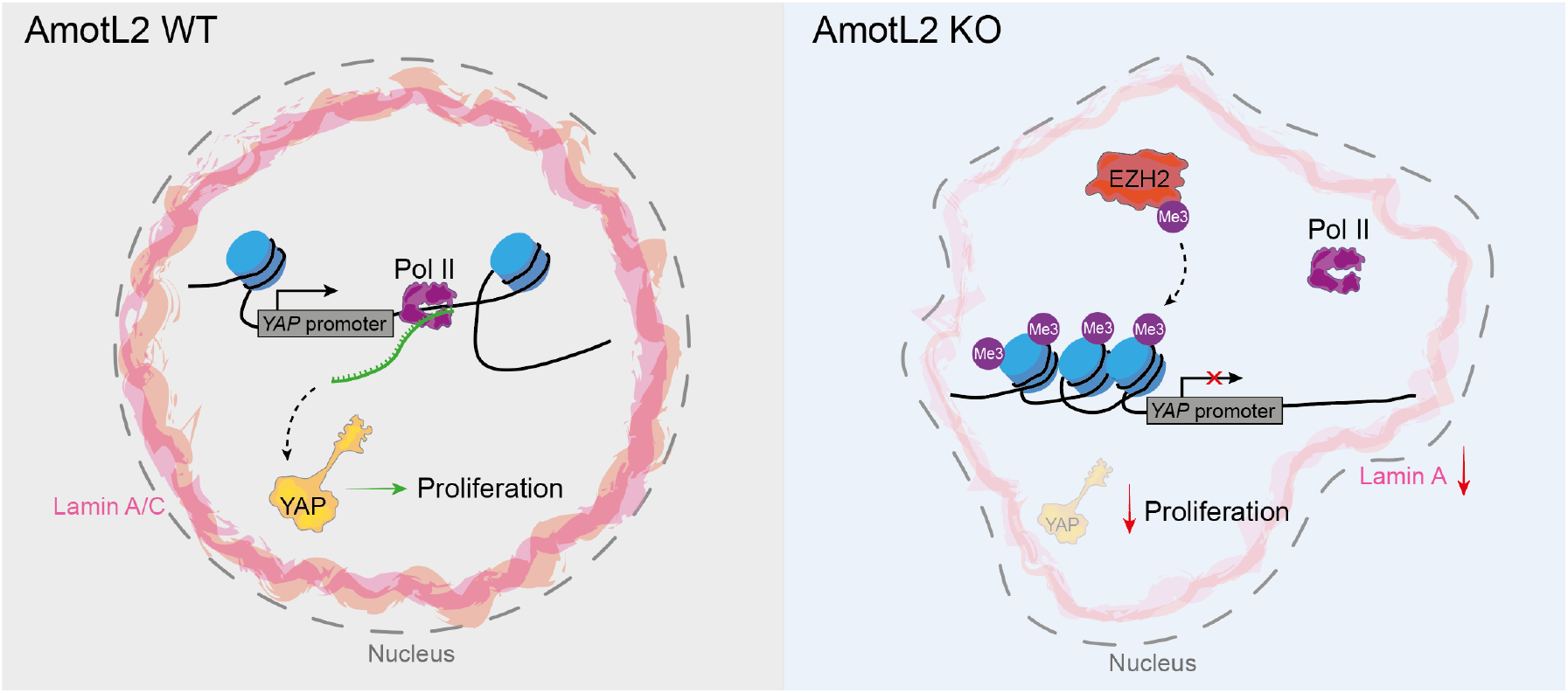
Model of AmotL2 dependent regulation of YAP. In AmotL2 WT endothelial cells, normal nuclear morphology and lamin A expression levels are observed, along with open chromatin and Pol II binding driving *YAP* transcription with concomitant proliferation. Conversely, deletion of endothelial AmotL2 leads to impaired nuclear lamin A levels, perturbed nuclear shape, changes to global chromatin conformation and increased H3K27me3 within the *YAP* promotor. This leads to reduced Pol II binding to the Y*AP* promotor, reduced *YAP* transcription and consequently, reduced proliferative capacity. Schematic drawn using Adobe Illustrator (v25.4.1).

## Materials and Methods

### Human AAA samples

Aortic tissue samples from patients with aortic abdominal aneurysm (AAA) were obtained from the surgery performed at Karolinska Hospital in Stockholm. Samples of the abdominal aorta of beating-heart, solid organ transplant donors were used as controls. Ethical permit (2009/1098-32) was granted by Regionala etikprövningsnämnden (Regional ethical review boards) in Stockholm. RNA extraction was performed from the medial layer of the aorta and subsequently sequenced by HTA 2.0 (Human Transcriptome Array 2.0 - Affymetrix) platform.

### Animals

For Yap/Taz iΔEC, *Wwtr1 flox/flox*; *Yap flox/flox* mice (Jackson Laboratory) were crossed to *Cdh5(BAC)-CreERT2*^51^ transgenic mice. Mice were used in mixed background or after backcrossing to C57BL/6 according to published protocol (Jackson Laboratory). Yap/Taz WT mice refers to *Wwtr1 flox/flox; Yap flox/flox* mice only. To induce endothelial-specific Yap/Taz gene inactivation, tamoxifen (Sigma, T5648) in corn oil (Sigma, C8267) was administered by oral gavage for 5 continuous days in 8-week-old mice (2 mg/mouse/day). *Cre*-positive tamoxifen-treated *Wwtr1 flox/flox; Yap flox/flox* mice were compared to control mice (Ctrl) that were tamoxifen-treated *Cre*-negative *fl/fl*. Analysis of aortic samples was performed 7-9 weeks after tamoxifen induced deletion. All animal experiments were approved by the National Animal Experiment Board in Finland. Ethical permits: ESAVI/15852/2022 and ESAVI/4975/2019.

For AmotL2 iΔEC, *amotl2 flox/flox* mice with loxP-flanked amotl2 gene, were crossed with *Cdh5(PAC)-CreERT2*^52^ and ROSA26-EYFP transgenic mice. To induce endothelial-specific amotl2 deletion, tamoxifen was administered by intraperitoneal (IP) injection for 5 continuous days. For adult mice over 6 weeks old, 100µl of tamoxifen (20mg/ml) was administered and analysis of aortic samples was performed four weeks following injections. All mice in this study were of the C57BL/6 background and included both female and male mice. Ethical permits were obtained from North Stockholm Animal Ethical Committee and all experiments were carried out in accordance with the guidelines of the Swedish Board of Agriculture. Ethical permits: (N129/15, 12931-2020 and 22902-2021).

### Cell culture

Human umbilical vein endothelial cells (HUVEC, Promocell, Heidelberg, Germany) were cultured in Endothelial cell medium (ECM; ScienCell) supplemented with 5% foetal calf serum (FCS) (v/v), 1% endothelial cell growth supplement (ECGS) and 1% Pen/Strep antibiotic cocktail and were cultured on 0.2% gelatin (Sigma-Aldrich) (w/v in PBS)-coated plates. Human umbilical arterial endothelial cells (HUAEC, Promocell, Heidelberg, Germany)) were cultured as above in endothelial cell basal medium (ECBM) MV medium (Promocell) supplemented with 2% foetal calf serum (FCS) (v/v), 0.4% Endothelial Cell Growth Supplement, 0.1 ng/ml epidermal growth factor (recombinant human), 1 ng/ml basic fibroblast growth factor (recombinant human), 90 μg/ml heparin and 1 μg/ml hydrocortisone. Both HUVEC and HUAEC were grown to passage 4 or 5.

### siRNA transfection

HUVEC and HUAEC were transfected with SMARTpool siRNA (Dharmacon) targeting YAP, TAZ or EZH2 in parallel with a scrambled (scr) control siRNA. Transfections were performed using Lipofectamine RNAiMax (Invitrogen) transfection agent and Opti-MEM I Reduced Serum Medium, GlutaMAX Supplement (Gibco) according to manufacturer’s instructions. Cells were transfected with 15 pmol siRNA in a twelve-well plate (1×10^5^ cells/well) and scaled accordingly. Briefly, for a single well of a twelve-well plate, 15 pmol siRNA duplexes were made in 125 μl of OptiMEM medium and incubated at room temperature for 5 min. For co-depletion of multiple targets (YAP and TAZ), 7.5 pmol per target (total 15 pmol) of siRNA duplexes was used. At the same time, 2 μl Lipofectamine was made up in 125 μl of OptiMEM and incubated at room temperature for 5 min. siRNA and Lipofectamine mix were combined and gently mixed before incubating at room temperature for 20 min. Following this, culture medium of cells seeded out the previous day, at a density 1×10^5^ cells, was aspirated and replaced with 500 μl of OptiMEM. Transfection mix was gently added to the cells dropwise. Cells were incubated in this mixture for 4-6 h, before transfection medium was aspirated and replaced with supplemented normal culture medium. siRNA-treated cells were incubated for 24-72 h before being used in downstream assays. The siRNA molecules used (from Dharmacon/Horizon Discovery) are shown below. ON-TARGETplus Human EZH2 (2146) siRNA – SMARTpool (5’-GAGGACGGCUUCCCAAUAA-3’, (5’-GCUGAAGCCUCAAUGUUUA-3’, (5’-UAACGGUGAUCACAGGAUA-3’, (5’-GCAAAUUCUCGGUGUCAAA-3’); ON-TARGETplus Human WWTR1 (25937) siRNA – SMARTpool (5’-CCGCAGGGCUCAUGAGUAU-3’, 5’-GGACAAACACCCAUGAACA-3’, (5’-AGGAACAAACGUUGACUUA-3’, (5’-CCAAAUCUCGUGAUGAAUC-3’); ON-TARGETplus Human YAP1 (10413) siRNA – Individual siRNA (5’-GCACCUAUCACUCUCGAGA-3’.

### Lentivirus generation

Lentiviral supernatants were produced in HEK293T cells following transfection using Lipofectamine 3000 (Invitrogen) transfection agent with 3^rd^ generation packaging plasmids and either shScr (SHC332; Sigma), shYAP (addgene #42541), FUW-tetO-wtYAP (addgene #84009) and FUW-tetO-HA-YAP5SA generously gifted by Stefano Piccolo, shAmotL2 (MXS19; Sigma), shAmotL2 #1 (GAACAAGATGGACAGTGAAAT) or shAmotL2 #2 (GAGAGATTGGAATCTGCAAAT) (VectorBuilder). HUVEC or HUAEC cells were transfected with the above lentiviral supernatants diluted (2:1) with ECMB supplemented media with polybrene (8μg/ml) (Sigma) overnight. The following day viral supernatants were removed and replaced with fresh supplemented media and cultured for 72-96h post transduction before downstream analysis.

### RT-PCR

Total RNA was isolated from HUVEC and HUAEC using RNeasy Plus Kit according to manufacturer’s instructions (Qiagen). Harvested mRNA was eluted from RNeasy spin columns in RNase free water and stored at −80°C. mRNA samples were quantified, and purity checked using the NanoDrop Lite Spectrophotometer (Thermo Scientific). Using High-Capacity RNA-to-cDNA Kit (Thermo Scientific) cDNA was synthesized according to the manufacturer’s instructions using 500-1000 ng RNA. Master mix and mRNA were combined in 9:11 respectively. Samples were placed into C1000 Thermal Cycler (Biorad) to incubate at 42°C for 60 min followed by 70°C for 5 min.

Gene expression of targets listed in supplemental table 2 were analysed by SYBR green-based qPCR analysis using Applied Biosystems QuantStudio 7 Real-Time PCR Systems (Thermo Scientific), following the standard PCR cycling sequence using *GAPDH* as an internal control. Results for qPCR target genes were normalised to internal *GAPDH* controls. Statistical analysis of qPCR data was carried out using ΔΔCt method.

### Western blotting and subcellular fractionation

Protein expression was analysed using precast Bis-Tris-polyacrylamide gel electrophoresis (PAGE)(Invitrogen). Cells were washed in 1x PBS before being lysed in 1x RIPA lysis buffer (EMD Millipore corp). Cell lysates were boiled at 100°C for 5 min, and mixed with NuPAGE LDS sample buffer (4x) (Invitrogen), NuPAGE Sample reducing agent 10x (Invitrogen) and total protein separated on a NuPAGE 4-12% Bis-Tris polyacrylamide gradient gel (Invitrogen) prior to western blotting on Pure Nitorcellulose 0.2 μm membranes (PerkinElmer). Membranes were probed with primary antibodies, followed by horseradish peroxidase (HRP) conjugated secondary ECL anti-mouse (NA931V; 17438421) or ECL anti-rabbit antibodies (NA934V; 17434242) (Amersham) were used for detection of probed proteins using enhanced chemiluminescence (ECL)(Perkin Elmer). Chemiluminescence was detected using an iBright FL1500 (ThermoFisherScientific) imager. Antibodies used for western blotting: Rabbit pAb anti-AmotL2 (affinity-purified as previously described^53^ (Innovagen, Lund, Sweden). Anti-YAP (D8H1X; #14074) XP® Rabbit mAb, rabbit mAb anti-pYAP Ser127 (also cross reacts with pTAZ Ser89)(D9W2I; #13008), rabbit mAb anti-TAZ (E8E9G; #83669) were from Cell Signalling Technologies. Mouse anti-GAPDH (ab181602) was from Abcam. Mouse anti-Lamin A/C (sc-7292; 636), anti-YAP/TAZ (sc-101199; 63.7) were from Santa Cruz Biotechnology. **Subcellular fractionation** for western blot analysis was performed using the NE-PER™ Nuclear and Cytoplasmic Extraction Reagents (Thermoscienctific – WC319307) kit according to manufacturer’s instructions.

### ChIP-Atlas

Publicly available ChIP-seq data submitted to the NCBI SRA was accessed from the ChIP-Atlas^54^. Predicted target genes of YAP were also devised using the ChIP-Atlas database. The following datasets were used for YAP1-ChIPseq: MCF7 (GSM3577938), (GSM3732639). (GSM2859575), (GSM3732641). (GSM3577954), T47D (GSM3577945), HEK293 TEAD1-ChIPseq: MCF7 T47D (GSM2859586), HEK293 TEAD4-ChIPseq: MCF7 T47D (GSM3577942), HEK293 (GSM3732642). Data were visualised using the IGV browser version 2.9.4. ENCODE ChIP-seq data used to view various histone marks around the YAP promotor included: HUVEC Pol II GSM1305213, HUVEC H3K27ac Accession: ENCFF656TFQ Experiment: ENCSR000ALB, HUVEC H3K4me3 Accession: ENCFF925RAZ Experiment: ENCSR000AKN, HUVEC EZH2 Accession: ENCFF700IMN Experiment: ENCSR000ATA, HUVEC H3K27me3 Accession: ENCFF938EXZ Experiment: ENCSR000AKK, and were visualised using the IGV-Web app version 1.12.5, igv.js version 2.13.5.

### Whole mount staining

Whole mount staining was performed as previously described^5^. Briefly, mice were euthanised and aorta harvested. The chest cavity was pinned open and piercing of left ventricle, whole body perfusion using 1x PBS (or in one cohort 1% PFA) was performed to clear the aorta of blood. Fat and surrounding adventitial tissue were dissected away and whole aortae further fixed in 4% PFA for 1 h at room temperature. Tissue was then cut length ways and for the aortic arch dissected as shown in figure Fig5e. Tissue was then mounted to wax blocks by pinning and tissue covered with a drop of PBS to ensure tissue doesn’t dry out while other pieces were mounted. Tissue was then fixed for another 10 min once mounted and washed with PBS. Tissue was permeabilised for 20 mins at RT using 0.1% Triton X-100 in PBS followed by 3x PBS washes at intervals of 15 min. Tissue was subsequently blocked in 5% FCS for 1 h at RT or overnight at 4°C. Primary antibody incubation was performed in blocking buffer with the following antibodies overnight at 4°C. Tissues was then washed 3x 30 min intervals with 1x PBS followed by 2ndary antibody incubation for 4-6 h at RT. Tissue were washed before mounting between two coverslips using Fluoroshield with DAPI (Sigma; F6057). Tissue was then imaged using either Zeiss 700, or Leica Thunder microscope. Images were taken with the same microscope settings for all samples within the same experiment as outlined under the image analysis methods. The following antibodies were used for immunofluorescent staining: Rabbit pAb anti-AmotL2 (affinity-purified as previously described^53^ (Innovagen, Lund, Sweden)), anti-YAP (D8H1X; #14074) XP® Rabbit mAb (CellSignallingTechnologies), mouse anti-YAP/TAZ (sc-101199; 63.7) (Santa Cruz Biotechnology), rabbit pAb anti-ki67 (ab15580), rabbit pAb anti-VE-cadherin (ab33168), EYFP was stained for using chicken pAb anti-GFP (ab13970) or goat pAb anti-GFP (ab6673), and rabbit anti-ERG mAb (ab92513) were from Abcam. TexasRed phalloidin (T7471) (Invitrogen) and phalloidin-Atto 647N (65906; Sigma) were used to label the actin cytoskeleton. Rat anti-Cd31 (MEC 13.3; 553370) and Rat anti-Cd144/VEcadherin (1104.1; 555289) were from BD Biosciences.

### Proximity ligation assay

Tissue was fixed and processed as described for immunofluorescence whole mount staining. Tissue was pinned out, permeabilised and washed a previously described. Tissue was then blocked as per the manufacturers conditions before proceeding with the assay using the Duolink In situ proximity ligation assay PLA kit (Sigma). Antibodies used for PLA; Anti-TEAD1 mouse mAb (610923) BD Biosciences, anti-YAP (D8H1X; #14074) XP® Rabbit mAb (CellSignallingTechnologies).

### Chromatin immunoprecipitation

HUVEC and HUAEC cells that were subjected to ChIP were transduced with lentiviral vectors for 72-96 h before being processed using iDeal ChIP-seq kit for Transcription factors (Diagenode), according to the manufacturers protocol with modifications. Briefly, cells were crosslinked with 1 % formaldehyde at RT for 15 min with gentle shaking. Glycine stop was added to the cells at 1:10 according to the protocol for 5 min before removal of medium and washing cells once with ice cold 1x PBS keeping cells on ice. Cells were then lysed in 15ml lysis buffer iL1 provided by the kit and incubated at 4 °C for 20 min with rotation. After centrifugation at 500 *g* for 5 min at 4°C, cell pellets were further lysed under rotation for 10 min in Lysis buffer iL2. Chromatin was sheared using the Covaris M220 focused ultrasonicator for 15 min and stored at −80 °C. For immunoprecipitation, ChIP grade antibodies were incubated with prewashed Protein A-coated magnetic beads for 2 - 4 h at 4 °C under rotation. Sheared chromatin was then added to beads and incubated overnight at 4°C under rotation. Beads were spun and washed according to the protocol. IP and input DNA were then eluted and purified according to the manufacturers protocol. Real-time PCR was carried out using 1% of IP or input DNA per reaction using SYBR green reagents according to the manufacturers protocol. Fold enrichment (2^-DDCT^) was calculated using input controls and IgG controls as background. Antibodies used for ChIP experiments were as follows: YAP (D8H1X) XP® Rabbit mAb (14074) (Cell signalling technologies), RNA pol II antibody (mAb) (39097), Histone H3K27me3 antibody (pAb) (39155) were from Active Motif. Mouse IgG (I8765) was from Sigma. Rabbit IgG (C15410206) (Diagenode). Anti-Histone H3 (acetyl K27) (ab4729) and anti-Histone H3 (tri methyl K4) (ab8580) were from Abcam. Primers for ChIP RT-qPCR were as follows: *AmotL2*; forward 5’-TGCCAGGAATGTGAGAGTTTC-3’, reverse 5’-AGGAGGGAGCGGGAGAAG-3’. *CTGF*; forward 5’-GAGCTGAATGGAGTCCTACACA-3’, reverse 5’-GGAGGAATGCTGAGTGTCAAG-3’. *YAP* (promotor region 1); forward 5’-GCCGTCATGAACCCCAAGA-3’, reverse 5’-GAGAGGGGCAACGAGGTTAC-3’. *YAP* (promotor region two – supplemental figure 10b-c); forward 5’-GCCTCTCGGTCCACTTCAG-3’, reverse 5’-GGTTCTACCCCCACCTCTAAT-3’.

### Incucyte growth curves

Growth curve analyses of HUVEC transfected with shScr or shAmotl2 lentivirus or treated with verteporfin (0.2μg/ml) or DMSO control were performed using Incucyte-Zoom live-cell microscope (Essen Bioscience). Twelve-well plate (1×10^5^ cells/well) were seeded to a 12 well plate 48h post-transfection. Cell coverage/confluence was calculated using built in Incucyte image analysis with masking of cell confluence which was calculated as a percentage of the maximum value obtained from the end point of the assay which was indicated when 100% confluency for control conditions was reached. 4 regions per well were imaged at hourly intervals for 65 h using the 20x objective.

### EdU proliferation assay

EdU incorporation was performed according to the manufacturers protocol using the Click-iT EdU Alexa Fluor 555 Imaging kit (Invitrogen) Lot: 2263481. Briefly, HUVEC were plated to Ibidi chamber slides and grown before being subjected to lentiviral transduction for 72 h before analysis of proliferation using the Click-iT EdU Cell proliferation kit for imaging Alexa fluor 647. Cells were then replated to 0.2% gelatin coated Ibidi chamber slides and left overnight to adhere and equilibrate. The following day, cells were incubated with EdU component A for 2 h at 37°C before fixing in 4% PFA and subsequent permeabilization, blocking and staining as dictated by the manufactures protocol. Cells were counterstained with Hoechst (@1:1000) and imaged using the Leica Thunder system. 3-5 images per condition were taken using 20x air objective using the Leica Thunder microscope.

### Hydrogels

Petrisoft™ easy coat hydrogel 0.2 kPa or 50 kPa 6-well plates were coated with 5μg/ml human fibronectin (Sigma) and RNA lysates harvested as described.

### DNA methylation sequencing

HUAEC cells were subjected to lentiviral knockdown of shScr or shAmotL2 before DNA isolation using the DNeasy Blood&Tissue Kit (Qiagen). DNA was bisulphite converted according to the protocol of the EZ DNA Methylation-Gold™ Kit (Zymoresearch) and subsequently sent for DNA MethylationEPIC Infinium bead array (illumine) sequencing. Bioinformatic analysis was performed using the R package RnBeads 2.0.^55^

### ATAC-seq

ATAC-seq was performed as previously reported^56^,^57^,^58^. Using the ATAC-seq library preparation ATAC-seq library preparation kit (ActiveMotif) HUVEC and HUAEC cells were transduced with shScr and shAmotL2 for 72 h before cells were trypsinised and counted using Trypan blue. 100,000 cells were centrifuged, and nuclei was extracted using ice-cold ATAC-lysis buffer (according to the kit protocol; Active Motif) and the pellet resuspended in the Tn5 tagmentation master mix. The transposition reaction was incubated at 37°C for 30 min at 800 rpm using a thermomixer. Tagmented DNA was purified according to the protocol’s column purification step, followed by PCR amplification of tagmented DNA for 10 cylces using Illumina’s Nextera indexed adapters. After the PCR reaction, libraries were purified with the SPRI magnetic beads and library quality, and concentration was assessed with Qubit bioanalyzer. 4 independent experiments were prepared for ATAC-seq for both HUVEC and HUAEC, resulting in 16 libraries in total. Libraries were then sequenced with the Illumina Hiseq platform with a 150bp paired-end repeats protocol.

Nextera adaptor sequences were firstly trimmed from the reads using skewer (0.2.2). These reads were aligned to a reference genome using BWA, with standard parameters. These reads were then filtered for high quality (MAPQ ≥ 13), non-mitochondrial chromosome, and properly paired reads (longer than 18 nt).

All peak calling was performed with macs2 using ‘macs2 callpeak --nomodel --keepdup all --call-summits’. For simulations of peaks called per input read, aligned and de-duplicated BAM files were used without any additional filtering. KEGG enrichment analysis was performed using KOBAS (v3), with a p value cut off <0.05. distribution of peak over various functional areas is analysed by ChIPseeker software. Bigwig files displaying sequenced tracks were visualised using the browser Integrative Genomics Viewer (v.1.12.5). Data were overlaid with existing ChIP-seq data of RNAP II, H3K27ac, H3K4me3, EZH2 and H3K27me3, to give contextual chromatin features of the *YAP* promotor.

### Image analysis

Confocal images were acquired using a scanning laser confocal Zeiss LSM700 microscope. Settings used were as follows: standard laser configuration, objectives -c-ApoChromat 40x/1.2 W Korr M27 water immersion lens, line averaging 2, Z stack 5 slices at 0.5uM step, resolution (1,024 × 1,024) pinhole diameter: 1 airy unit. For whole mount aortic arches imaged using tiled scan, Leica Thunder LED based microscopy was used using a 20x air objective – higher resolution regions of the aortic arch where then imaged using scanning laser confocal Zeiss LSM700 microscopy using the settings outlined above. Microscope settings were kept constant across conditions of experiments and between different independent tissue samples within an experiment.

For YAP nuclear quantification, fluorescent intensity across the major of axis of the cell was measured using DAPI as a marker of the nuclear region and intensity profiles exported and averaged using GraphPad prism. Fluorescent intensity of 60 cells in total were analysed from the aorta and inferior vena cava of 3 individual mice. To analyse nuclear YAP:TEAD1 PLA signal, the ImageJ plugin *BioVoxxel>Speckle Inspector* was used. Images were subject to Max Intensity projection before thresholding and then analysis of dots/nuclei. Junctional staining and quantification were analysed in ImageJ by drawing a line across the image (3 lines per image) and quantifying the staining intensity profile using the ‘*plot profile*’ function. The x and y coordinates were exported, and average staining intensity calculated using the area under the curve in Graphpad prism. For YAP intensity profiles of YAP/TAZ iΔEC quantification, whole image YAP intensity was analysed.

For analysis of nuclear circularity in YAP/TAZ iΔEC mice, confocal images were analysed using the DAPI staining of the nucleus using ImageJ. Images were analysed using the MorphoLibJ plugin. *MorphoLibJ>Segmentation>Morphological segmentation*. Images were processed as follow: Z stacks were stacked with Max projection; background was subtracted using ‘subtract background’ with sliding paraboloid and rolling ball radius of 50 pixels. A threshold was applied to the image to obtain a masked image before the application of Gaussian blur (Sigma: 2). The plugin MorphoLibJ was then used to segment images based on morphology using the following settings: *Run MorphoLibJ>Segmentation>Morphological segmentation (Settings: (Object image), Gradient – Morphological, radius 1, Watershed Segmentation – tolerance 15, overlaid basins > create image)*. The Segmented image was then used to analyse nuclear parameters using *MorphoLibJ>label map* followed by *MorphoLibJ>Analyze tools*. Data were then imported to GraphPad Prism for statistical analysis.

### Statistical analysis

All statistical analyses were performed using GraphPad Prism (v9), with the exception of ATAC-seq datasets in which *P* values were generated with the software outlined in supplemental table 3 and human AAA samples, where the correlation between AmotL2:YAP and AmotL2:TAZ were analysed using Pearson correlation and Pearson correlation coefficient generated using R (https://www.r-project.org/index.html). Statistical tests used to analyse differences between groups are outlined in each figure legend and consist of, unpaired Student’s *t*-tests, Mann-Whitney, and one- or two-way analysis of variance (ANOVA) with correction for multiple comparisons (Tukey’s, Šídák’s, or Dunnet’s) as specified in the figure legends. Normal distribution was tested using D’Agostino-Pearson test and non-parametric tests applied when appropriate. All tests were two-tailed and unpaired unless specified otherwise. The number of independent biological replicates is indicated in the figure legends and is referred to as *n. P* values corresponding to symbols used in the figures are outlined in the figure legend and denoted: *p<0.0332, **p<0.0021, ***p<0.0002, ****p<0.0001. Data are displayed as means +/- standard deviation unless otherwise specified. Sample sizes were not predetermined using any tests. Experiments were not randomized or blinded in any way.

For image analysis, a minimum of three replicate images (from the same sample) were taken from at least 3 independent experiments. Representative images shown are of a minimum of 3 mice or 3 independent experiments for *in vitro* cell cultures.

## Acknowledgments

This study was supported by grants from the Swedish Heart and Lung Foundation (LH), Novo Nordisk Foundation (NNF15CC0018346) (LH), the Swedish Cancer Society (LH), the Swedish Childhood Cancer Foundation (LH), Cancer Society of Stockholm (LH), the Swedish Research Council (LH), Knut and Alice Wallenberg Foundation (LH), European Research Council (ERC) under the EU Horizon 2020 research and innovation programme (grant agreement 773076, PS), Sigrid Jusélius Foundation (PS), Cancer Foundation Finland (PS) and Academy of Finland (PS).

## Author contributions

The study was conceived and designed by AJM and LH. AJM and HZ performed *in vitro* experiments. AJM, YZ and YvW performed *in vivo* experiments. AJM, OB and HMB performed *in silico* and bioinformatic analyses. PS provided samples from Yap/Taz transgenics. AJM wrote the manuscript and all authors contributed to the drafting and editing of the manuscript.

---

**Supplementary Figure 1.**
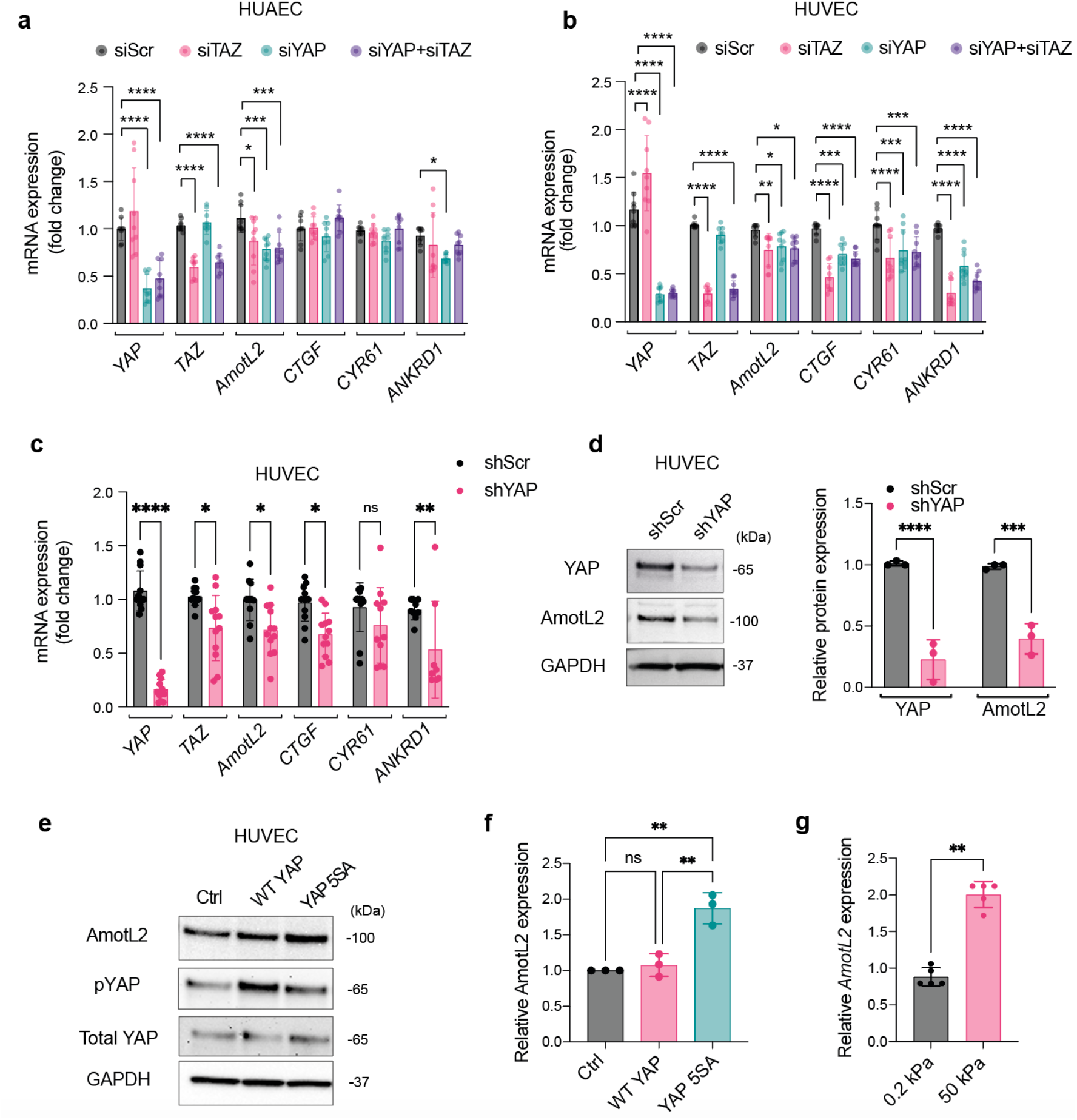
**a**, Fold change in mRNA expression of AmotL2 and known YAP target genes (*CTGF, CYR61* and *ANKRD1*) in HUAEC cells transfected with scrambled siRNA (siScr), siTAZ, siYAP or codepletion of YAP and TAZ (siYAP+siTAZ) analysed by qPCR and normalised to GAPDH. *n*=3 independent experiments, each with 3 technical replicates. (mean± s.d., 2way ANOVA with Dunnett’s multiple comparisons). **b**, As in panel a, in HUVEC *n*=3 independent experiments, each with 3 technical replicates. **c**, Fold change in mRNA expression of *AmotL2* and known YAP target genes (*CTGF, CYR61* and *ANKRD1*) in HUVEC cells transduced with shScr or shYAP lentivirus, analysed by qPCR and normalised to GAPDH. *n*=3 independent experiments, each with 3 technical replicates. (mean± s.d., 2way ANOVA with Dunnett’s multiple comparisons). **d**, Representative western blot showing YAP and AmotL2 expression in HUVEC cells transduced with shScr or shYAP lentivirus for 96h prior to immunoblot analysis. GAPDH was used as a loading control and normalisation for respective quantification shown in the right-hand panel, *n*=3 independent experiments. (mean± s.d., 2way ANOVA with Dunnett’s multiple comparisons). **e**, Representative western blot and quantification (**f**), showing AmotL2, total and phosphorylated YAP expression in none infected, WT YAP and YAP 5SA-overexpressing HUVEC, *n*=3 independent experiments. (mean± s.d., 2way ANOVA with Dunnett’s multiple comparisons). **g**, RT-qPCR of AmotL2 from HUVEC plated to 0.2 or 50 kPa hydrogels, (*n*=5 independent experiments, mean± s.d., Mann-Whitney).

**Supplementary Figure 2.**
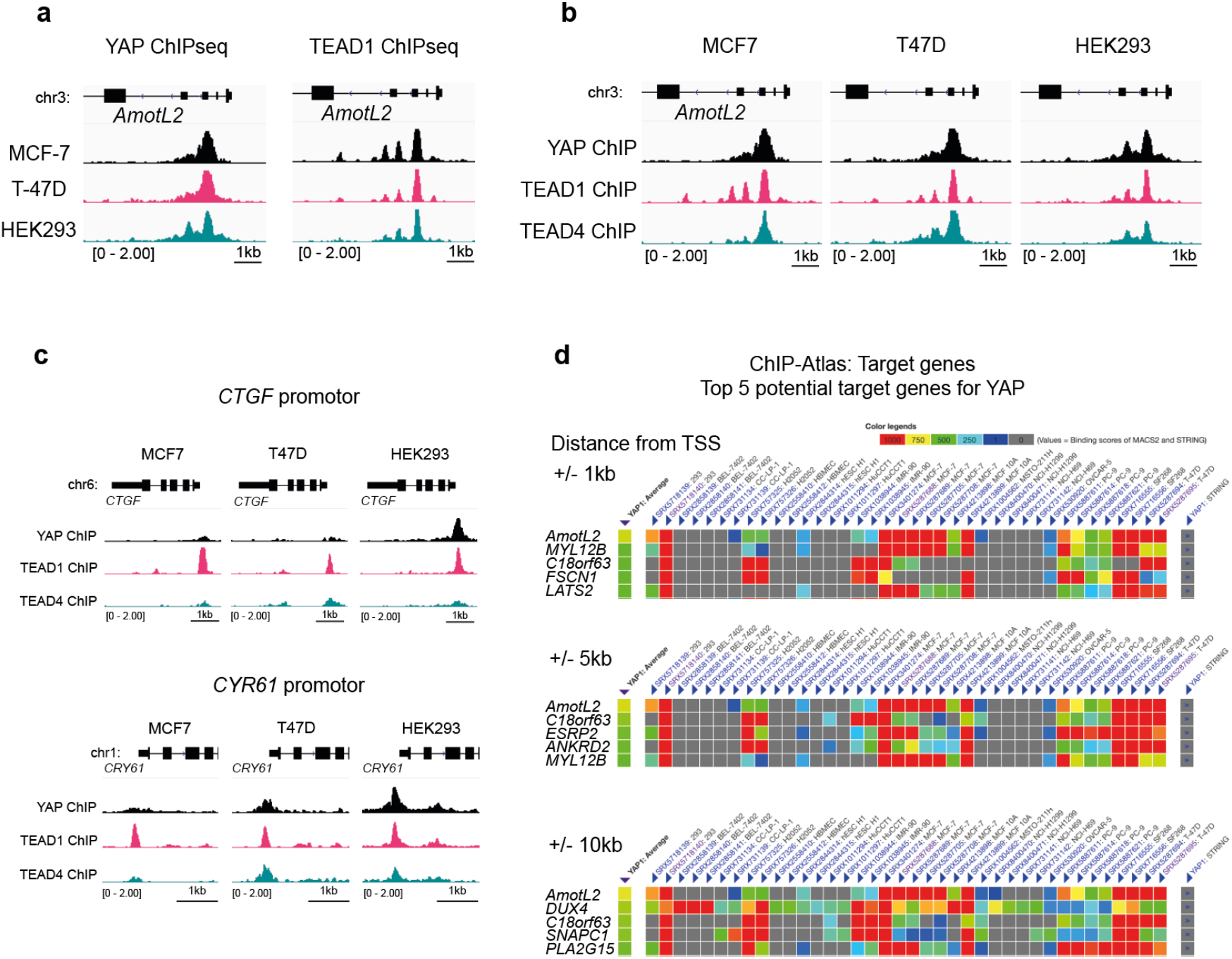
**a**, Genomic tracks displaying ChIP-Atlas (https://chip-atlas.org/) data of YAP (left panel) and TEAD1 (right panel) ChIP-seq data across MCF7, T47D, and HEK293, within the *AmotL2* promotor (Data sources are referenced in the methods). **b**, Genomic tracks displaying overlayed YAP, TEAD1 and TEAD4 ChIP-seq enrichment at the *AmotL2* promotor of indicated cell lines. **c**, Genomic tracks displaying overlayed YAP, TEAD1 and TEAD4 ChIP-seq enrichment at the *CTGF* and *CYR61* promotor of indicated cell lines. **d**, Top 5 hits of ChIP-Atlas predicted target genes bound by YAP in indicated datasets at 1, 5 and 10 kb from the transcriptional start site (TSS) of indicated target genes.

**Supplementary Figure 3.**
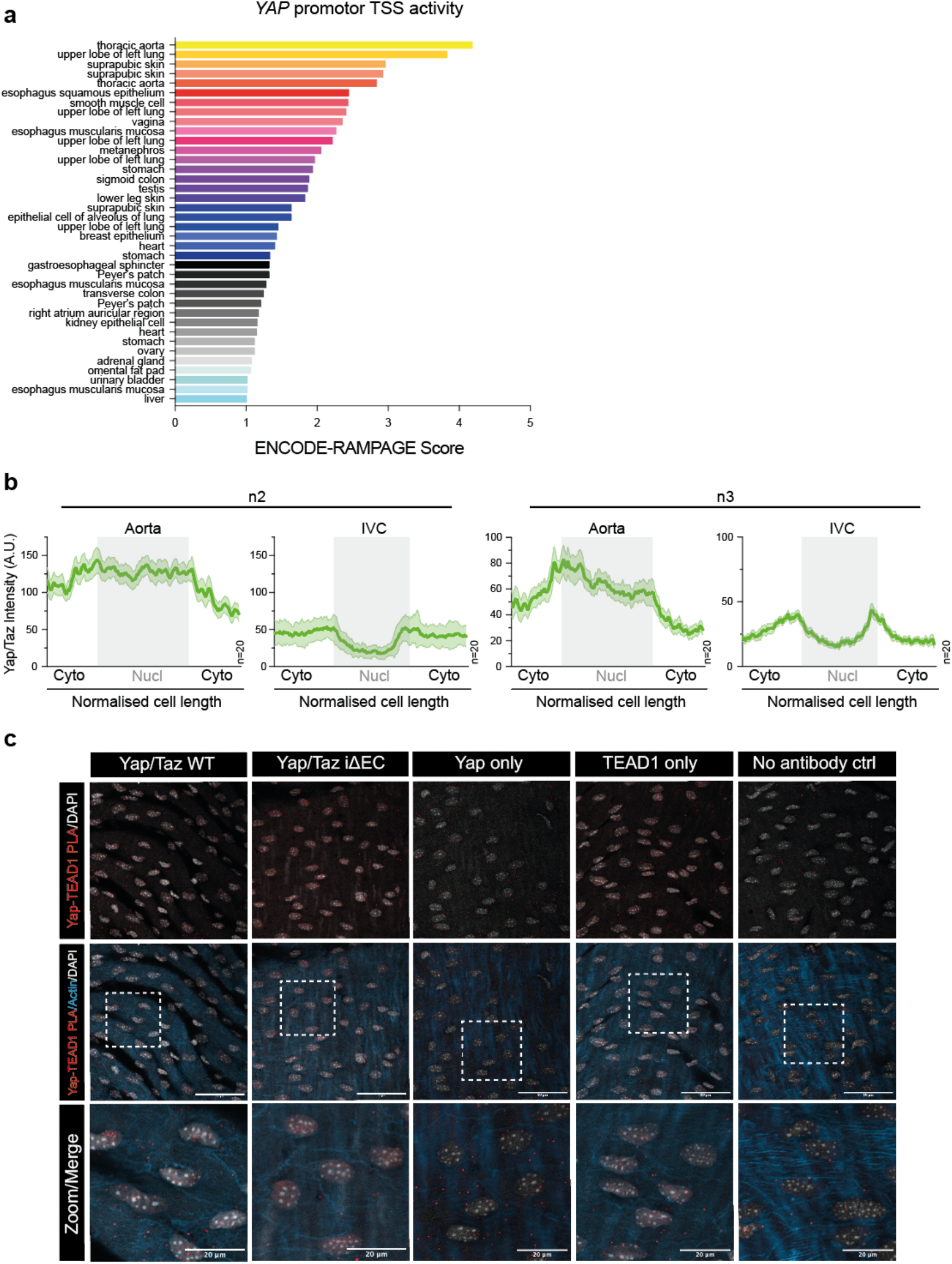
**a**, RNA Annotation and Mapping of Promoters for the analysis of Gene Expression (RAMPAGE) analysis of the transcriptional start sites (TSSs) of the YAP promotor, scored and ranked in descending order using SCREEN: Search Candidate cis-Regulatory Elements by ENCODE; https://screen.wenglab.org/.^59^ **b**, Quantification of nuclear and cytoplasmic staining of YAP/TAZ in the aorta and IVC. Histogram graph depicts the average intensity (mean± s.e.m.) of *n*=20 cells each from two independent animals labelled n2 and n3 (n1 is shown in Fig1d). A total of 60 cells per aorta or IVC were measured from *n*=3 mice. **c**, Representative images of *ex vivo en face* aortic tissue of both YAP/TAZ WT (*n*=5) and Yap/Taz iΔEC (*n*=5) inducible knockout mice, subjected to PLA showing the interaction between YAP and TEAD1 (indicated by red dots) and single antibody and no antibody controls. Quantification is shown in Fig2c. Nuclei are shown in grey and actin cytoskeleton in blue. Scale bar, 50µm and 20µm, for 40x and indicated zoomed regions, respectively.

**Supplementary Figure 4.**
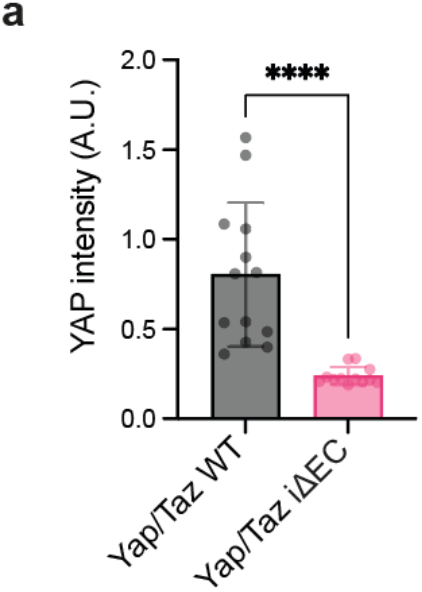
**a**, Bar graph indicates quantification of Yap fluorescent intensity, where each data point represents intensity profile from one image, 3 images/aorta (n=5 mice/group) were analysed, mean± s.d., Mann-Whitney).

**Supplementary Figure 5.**
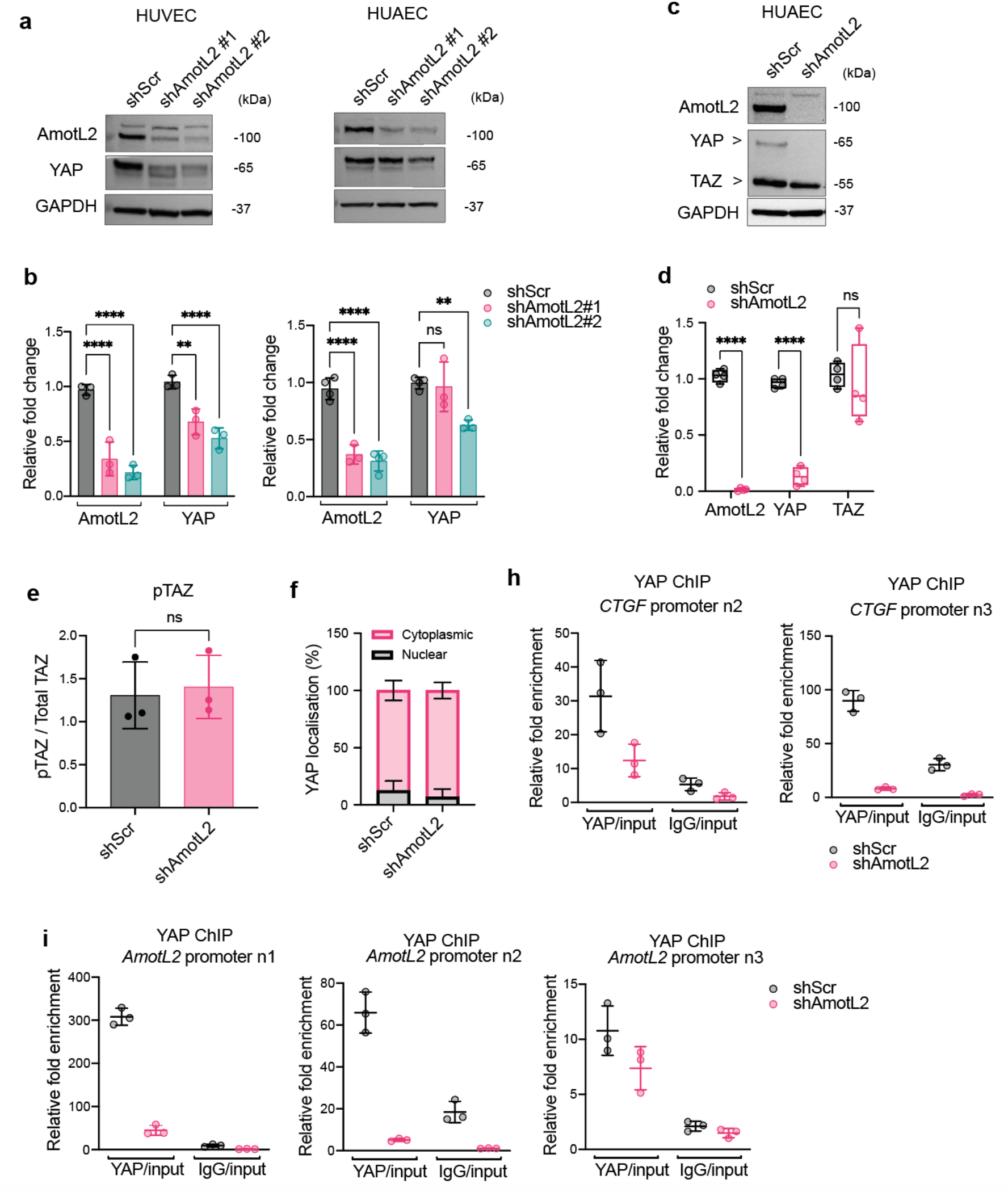
**a**, Western blot analysis of AmotL2 and YAP in HUVEC and HUAEC cells 96h post-treatment with lentivirus encoding shScr or two additional shRNA constructs targeting AmotL2 (shAmotL2#1 and shAmotL2#2). GAPDH was used as a loading control. Membranes are representative of n=3 independent experiments. **b**, Quantification of AmotL2 and YAP protein levels from panel a, relative to GAPDH loading control. *n*=3 independent experiments for both HUVEC and HUAEC, mean± s.d., 2way ANOVA with Dunnett’s multiple comparisons. **c**, Western blot analysis of shAmotL2 treated HUAEC using an antibody with specificity for both YAP and TAZ. Box plots shown in **d**, indicate quantification of AmotL2, YAP, and TAZ relative to GAPDH loading control, *n*=4, mean± s.d., 2way ANOVA with Dunnett’s multiple comparisons. **e**, Quantification of pTAZ ser89 levels, relative to total TAZ shown in Fig4e. (*n*=3 independent experiments, mean± s.d., Mann-Whitney). **f**, Quantification of nuclear:cytoplasmic fractionation and probing of YAP localisation, shown in Fig4g. HUVEC cells 96h post-treatment with shScr or shAmotL2 lentivirus. GAPDH and lamin A/C were used as positive and negative controls and were used for normalisation for quantification. *n=3* independent experiments. ChIP showing YAP binding to *CTGF* promotor (**h**,) and *AmotL2* promotor (**i**,) of shScr or shAmotL2 treated HUVEC. ChIP qPCR was performed using SBYR green reagents and quantification was normalised to an IgG control antibody. Plot shown is a representative experiment from n=3 independent experiments (Fig3h shows *n*=1 for the *CTGF* promotor). Each data point represents a technical repeat within one independent experiment (performed in triplicate). Graphs display (mean± s.d.).

**Supplementary Figure 6.**
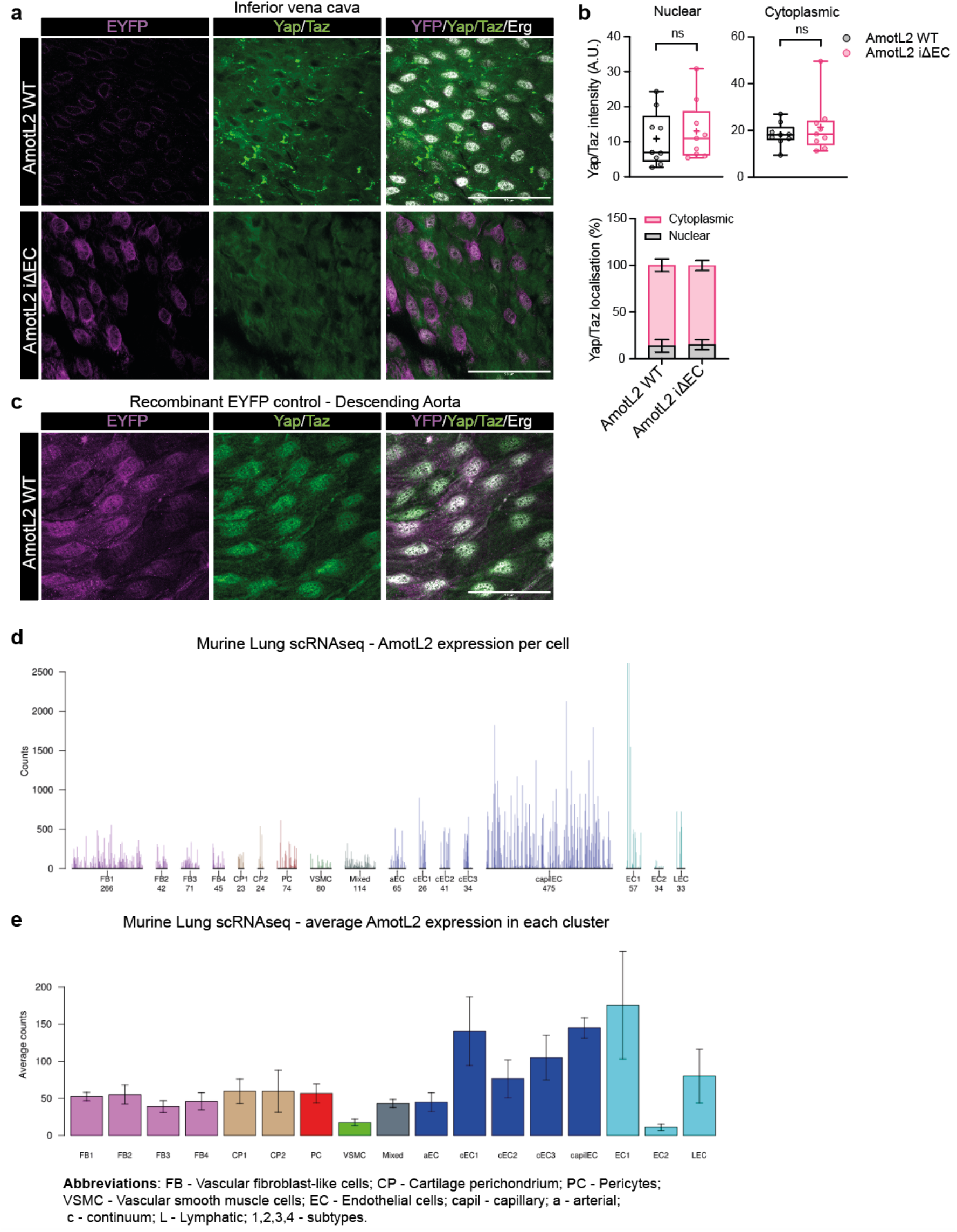
**a**, Representative images of *en face* staining of EYFP (magenta), Yap/Taz (green), and ERG (grey) in the inferior vena cava of both AmotL2 WT and AmotL2 iΔEC mice. Scale bar, 50µm. Images are representative of *n=3* mice/group. **b**, Quantification of Yap/Taz nuclear:cytoplasmic localisation and immunofluorescent intensity of the inferior vena cava of both AmotL2 WT (*n*=3) and AmotL2 iΔEC (*n*=3) mice. **c**, Staining as in a, of recombinant EYFP control murine aorta indicating that induction of EYFP expression in WT AmotL2 cells does not affect Yap/Taz expression. Scale bar, 50µm. Images are representative of *n=3* mice/group. **d**, Raw counts of AmotL2 expression across cell types of adult lung from scRNAseq data accessed through (https://betsholtzlab.org/VascularSingleCells/database.html)^60-24^ **e**, As in d, but displaying average counts.

**Supplementary Figure 7.**
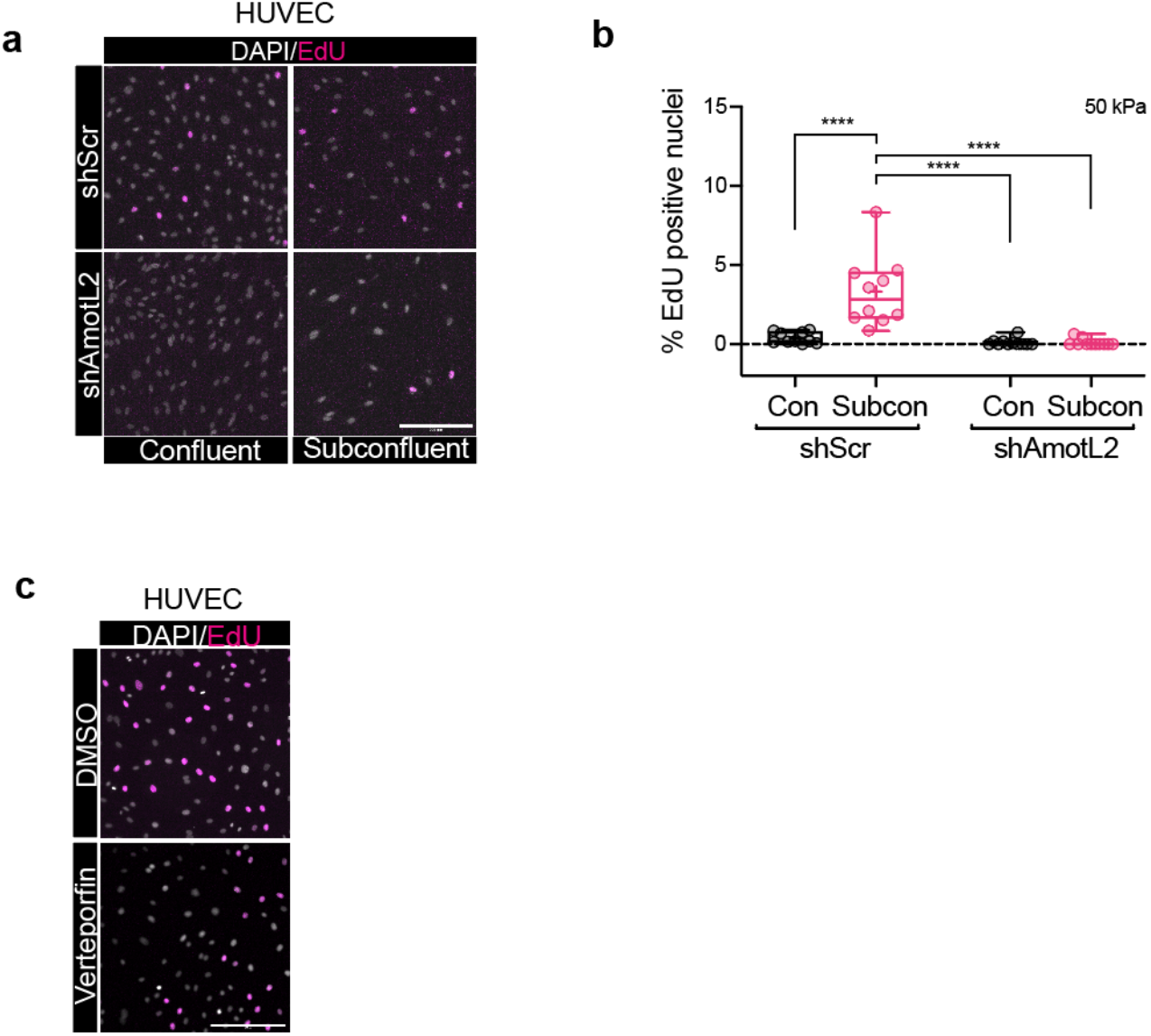
**a**, Representative images of shScr and shAmotL2 treated HUVEC 72h post infection, replated to gelatin coated plastic in confluent or subconfluent conditions. Incorporated EdU was detected with secondary antibodies and counterstained with Hoechst. Scale bar, 250µm. **b**, Box plots show quantification of EdU incorporation of HUVEC replated to gelatin coated 50kPa hydrogels following 48h post-lentiviral transduction with shScr or shAmotL2 lentivirus, where % of EdU positive cells was calculated against total number of cells stained with Hoechst. Each data point represents one field of view from *n*=3 independent experiments. (mean± s.d., 2way ANOVA with Dunnet’s multiple comparisons). **c**, Representative images of EdU positive HUVEC treated for 48h with 0.2µg/ml Verteporfin or DMSO vehicle. Scale bar, 50µm. Cells were counterstained with Hoechst. Scale bar, 50µm.

**Supplementary Figure 8.**
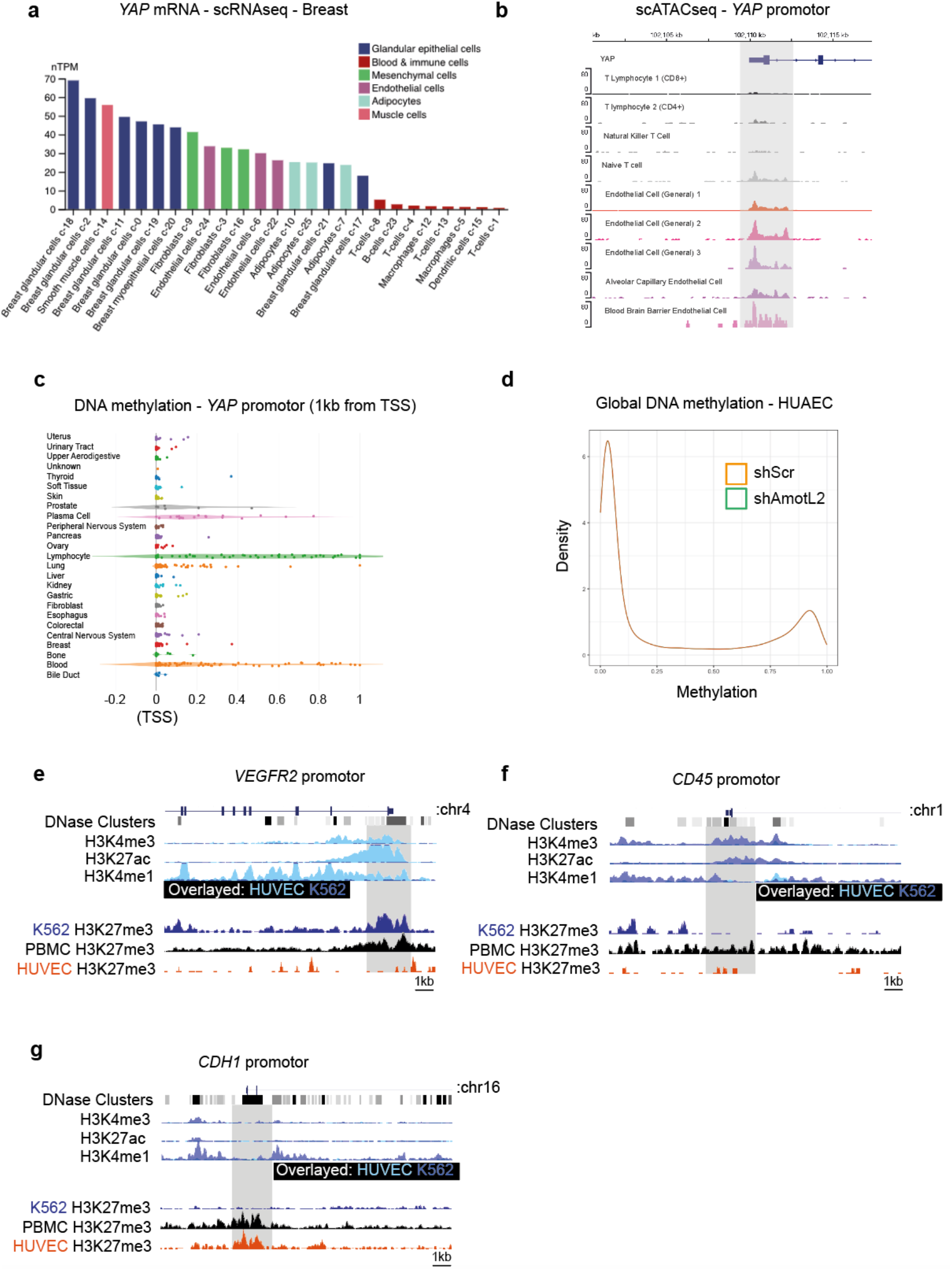
**a**, Screengrab of the Human protein atlas showing *YAP* mRNA expression from scRNAseq data of human breast cancer samples. **b**, scATACseq data indicating chromatin conformation around the *YAP* promotor using (http://catlas.org/catlas_hub/). Histograms indicate chromatin accessibility from specific cell types. **c**, publicly available DNA methylation data from the Dependency Map (Depmap.org) portal, showing DNA methylation of indicated cell types derived from human samples. **d**, Infinium EPIC array of 850,000 CpG sites across the genome. Data are averages of *n=4* for both shScr and shAmotL2 HUAEC samples. Histograms show the average distribution of DNA methylation profile of shScr (orange) and shAmotL2 (green). **e**, Screengrab of the UCSC browser displaying genomic tracks of ENCODE data of H3K4me3, H3K27ac and H3K4me1 ChIP-seq data across HUVEC and K562 (overlayed)(Data sources are referenced in the methods) and H3K27me3 of ChIP-seq data across K562, PBMC and HUVEC (Data sources are referenced in the methods), within the *VEGFR2* promotor (**e**), *CD45* promotor (**f**), *CHD1* promotor (**g**).

**Supplementary Figure 9.**
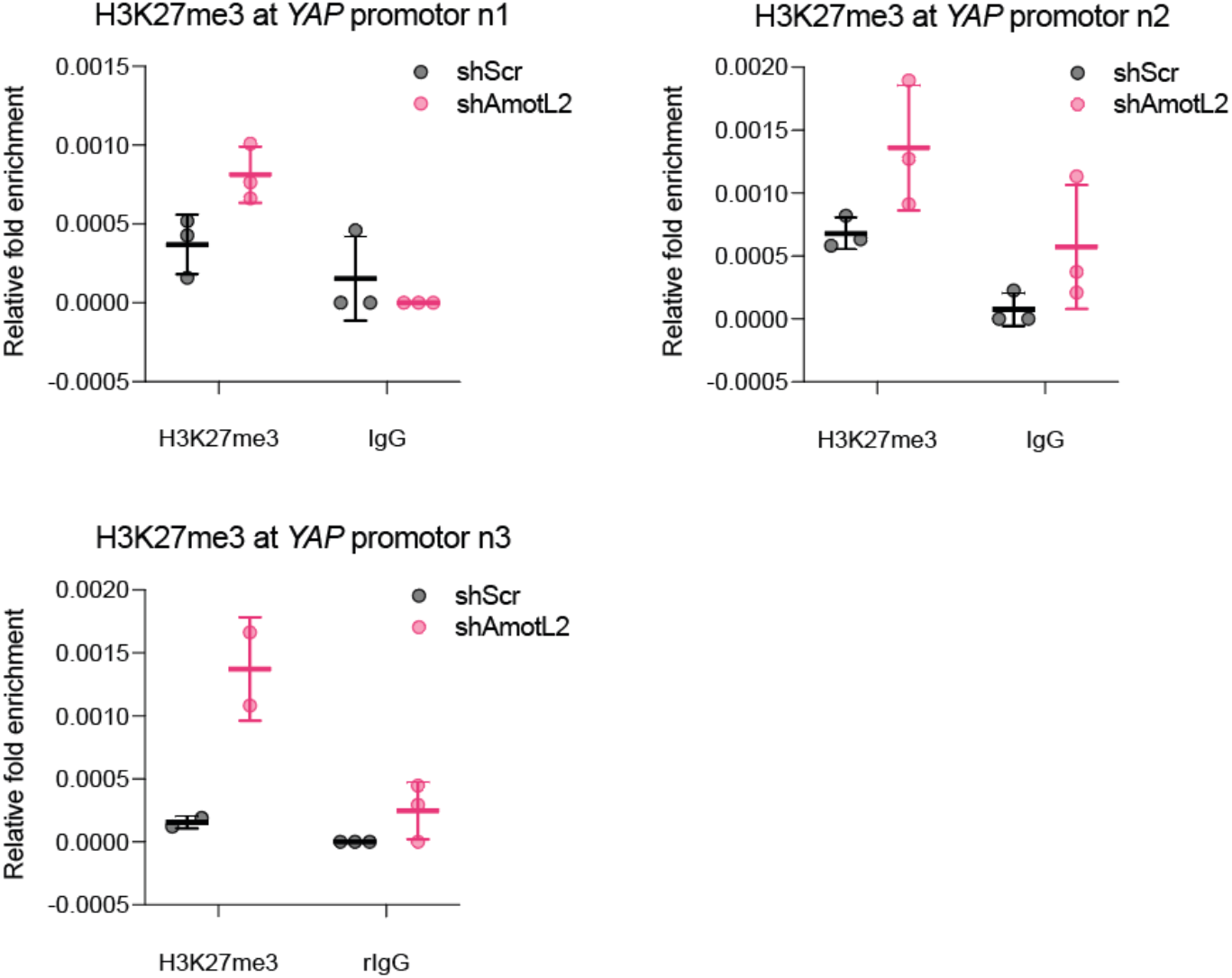
ChIP showing H3K27me3 pulldown at the *YAP* promotor of shScr or shAmotL2 treated HUAEC. ChIP qPCR was performed using SBYR green reagents and quantification was normalised to input and IgG control. Plots shown are representative of n=3 independent experiments. Each data point represents a technical repeat within one independent experiment (performed in triplicate). Graphs display (mean± s.d.).

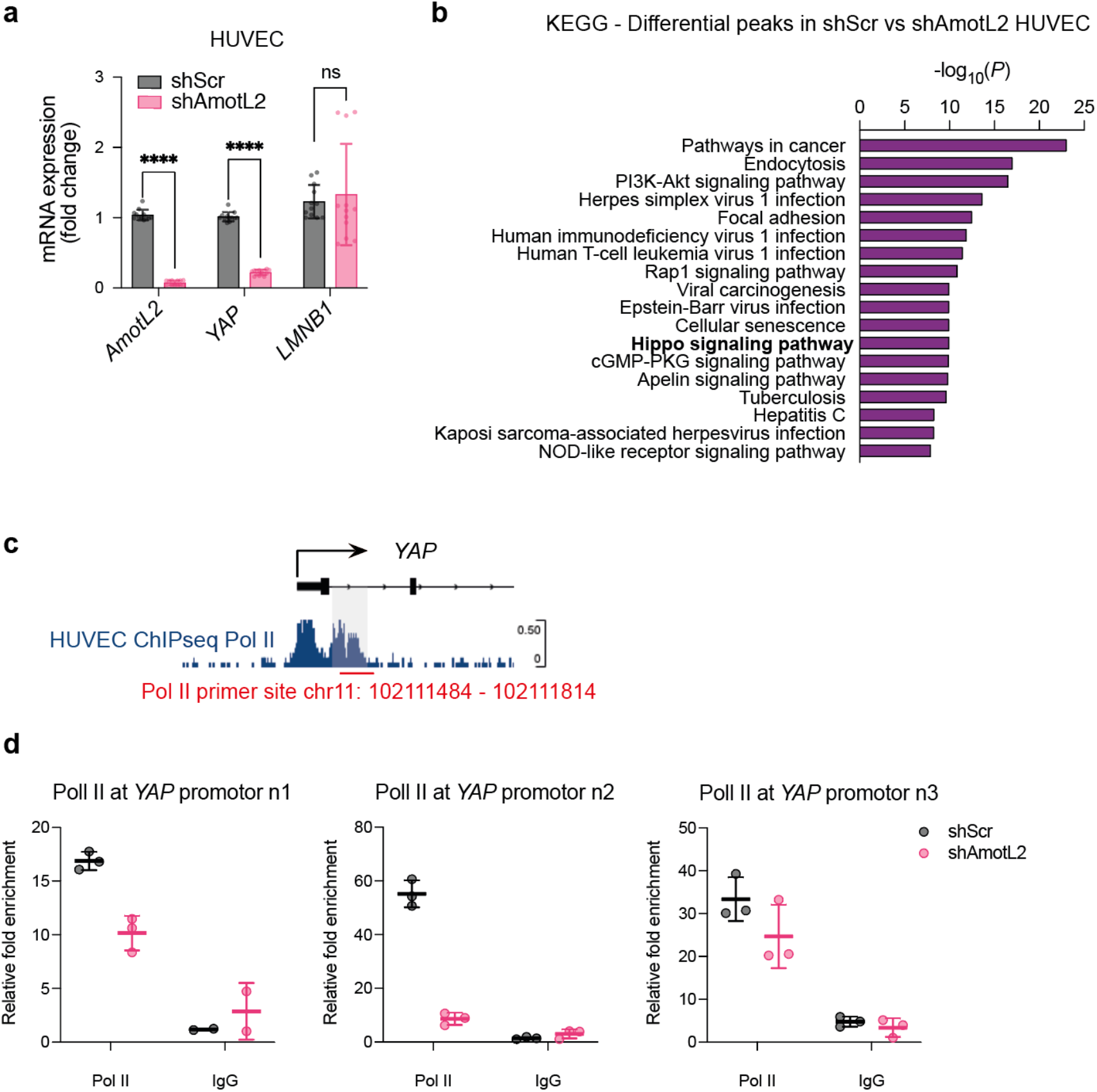
**a**, SYBR green RT-qPCR of *AmotL2, YAP*, and *LAMINB1* relative to housekeeping gene *GAPDH*, in AmotL2 knockdown HUVEC cells. (*n*=4 independent experiments, mean± s.d., 2way ANOVA with Šidák’s multiple comparisons). **b**, Top 20 differential KEGG pathway analysis of shScr, shAmotL2 transduced HUVEC indicating pathways with differential peak accessibility. Note ‘Hippo signalling pathway’ in bold. **c**, Schematic of primers designed for ChIP analysis of region highlighted in the second peak of bimodal accessibility within the *YAP* promotor where differential accessibility was observed from ATAC-seq data shown in Fig7i-j. **d**, ChIP showing Pol II pulldown at the region of the *YAP* promotor show in (b) of shScr or shAmotL2 treated HUVEC. ChIP qPCR was performed using SBYR green reagents and quantification was normalised to input and IgG control. Plots shown are representative of *n*=3 independent experiments. Each data point represents a technical repeat within one independent experiment (performed in triplicate). Graphs display (mean± s.d.).

**Supplemental table 2.**
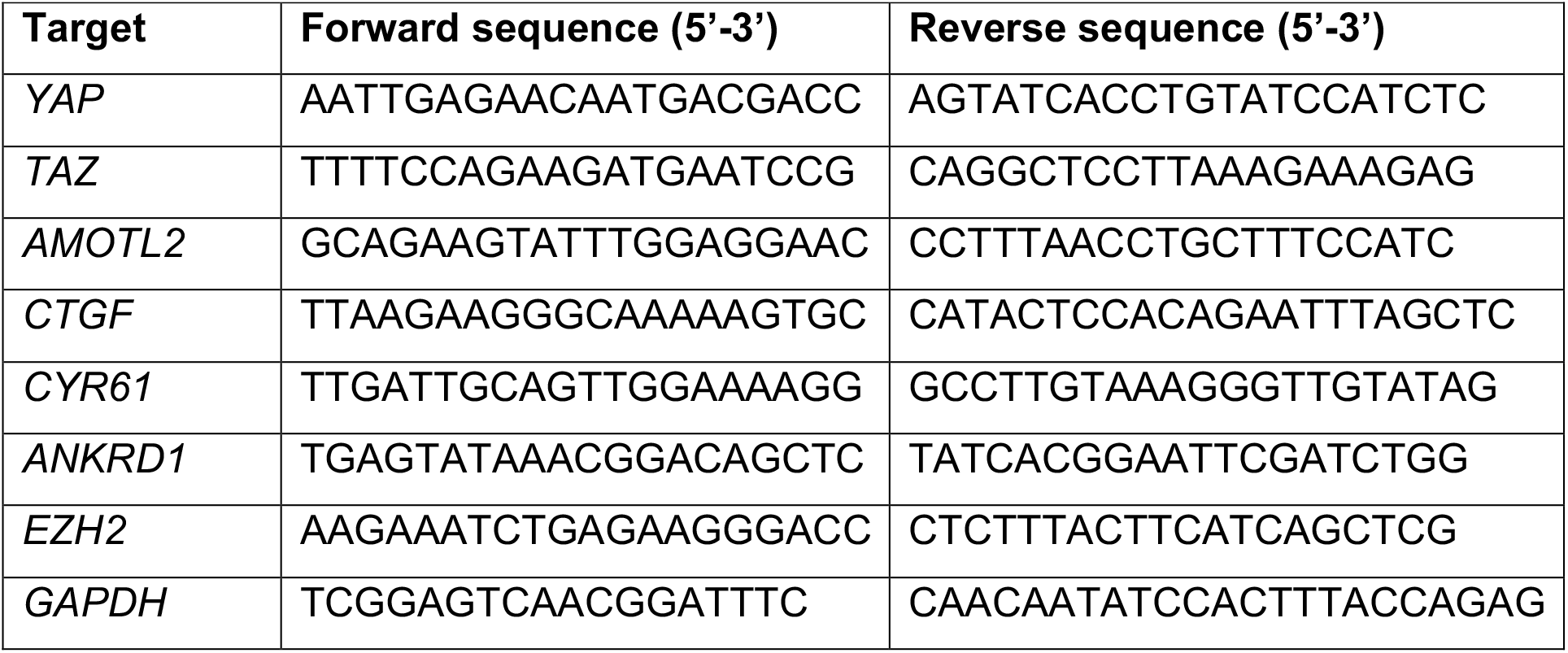

**Supplemental table 3.**

| Analysis | Software | Version | Parameters | Remarks |
| --- | --- | --- | --- | --- |
| Trimming | skewer | 0.2.2 | -m pe | Filter rawdata |
| QC | fastqc | v0.11.5 |  |  |
| mapping | BWA | 0.7.12-r1039 | -T 25 -k 18 | Mapped to the reference genome |
| correlation between samples | deepTools | 3.0.2 | --corMethod pearson |  |
| peak calling | MACS2 | 2.1.2 | -q 0.05 --call-summits --nomodel --shift -100 --extsize 200 --keep-dup all |  |
| Identification of motif | homer<br>findMotifsGenome.pl | v4.9.1 | -gc -len 8,10,12,14 |  |
| GO enrichment | Goseq, topGO, Bioconductor (2.13) | 4.10.2 | corrected pvalue<0.05 |  |
| KEGG enrichment | KOBAS | 3 | corrected pvalue<0.05 |  |

